# Deep mutational scanning of H5 hemagglutinin to inform influenza virus surveillance

**DOI:** 10.1101/2024.05.23.595634

**Authors:** Bernadeta Dadonaite, Jenny J. Ahn, Jordan T. Ort, Jin Yu, Colleen Furey, Annie Dosey, William W. Hannon, Amy L. Vincent Baker, Richard J. Webby, Neil P. King, Yan Liu, Scott E. Hensley, Thomas P. Peacock, Louise H. Moncla, Jesse D. Bloom

## Abstract

H5 influenza is a potential pandemic threat. Previous studies have identified molecular phenotypes of the viral hemagglutinin (HA) protein that contribute to pandemic risk, including cell entry, receptor preference, HA stability, and reduced neutralization by polyclonal sera. Here we use pseudovirus deep mutational scanning to measure how all mutations to a clade 2.3.4.4b H5 HA affect each phenotype. We identify mutations that allow HA to better bind α2-6-linked sialic acids, and show that some viruses already carry mutations that stabilize HA. We also identify recent viral strains with reduced neutralization to sera elicited by candidate vaccine virus. Overall, the systematic nature of deep mutational scanning combined with the safety of pseudoviruses enables comprehensive characterization of mutations to inform surveillance of H5 influenza.

## Main

Experimental research on H5 influenza and studies of past influenza pandemics have identified properties of several viral genes (e.g., HA (*1–3*), PB2 (*4*, *5*), and NP (*6–8*)) that increase the risk that animal influenza viruses pose to humans. Mutations in HA can increase pandemic risk by affecting several molecular phenotypes. One key phenotype is the ability of HA to use α2-6-linked rather than α2-3-linked sialic acid as its primary receptor (*9–11*), which is important since α2-6-linked sialic acids dominate the upper respiratory tract in humans (*12*). Another relevant phenotype is HA stability, as mutations that stabilize HA are frequently present in viruses capable of transmitting by the airborne route (*1–3*, *13*, *14*). HA’s antigenicity is another important phenotype: a component of pandemic preparedness is developing candidate strains for vaccine production (*15*), but new HA variants can contain mutations that reduce neutralization by antibodies elicited by mismatched vaccine strains. Finally, the ability of HA to mediate cell entry is an essential phenotype for viral fitness, so sites where mutations impair cell entry are good targets for HA-directed therapeutics (*16*, *17*).

While previous work has shown the importance of these HA phenotypes, the entire body of prior experimental research has only measured how these phenotypes are impacted by a small fraction of the many possible mutations to H5 HA. Therefore, it is often impossible to assess how newly observed HA mutations impact pandemic risk without performing experiments on each new mutation, an approach that lags behind virus evolution. Here we address this challenge by prospectively measuring how all possible amino-acid mutations to HA affect the phenotypes of cell entry, α2-6-linked sialic-acid usage, stability, and serum antibody neutralization (**Fig. 1A**). We make these measurements by applying a safe pseudovirus deep mutational scanning system (*18*) to the HA of a current WHO-recommended 2.3.4.4b clade H5 candidate vaccine virus (*15*) (**Fig. 1B-C**). The resulting maps of mutational effects enable real-time interpretation of the phenotypic effects of HA mutations observed during viral surveillance, including the ongoing H5N1 outbreak in dairy cattle (*19*).

**Figure 1.**
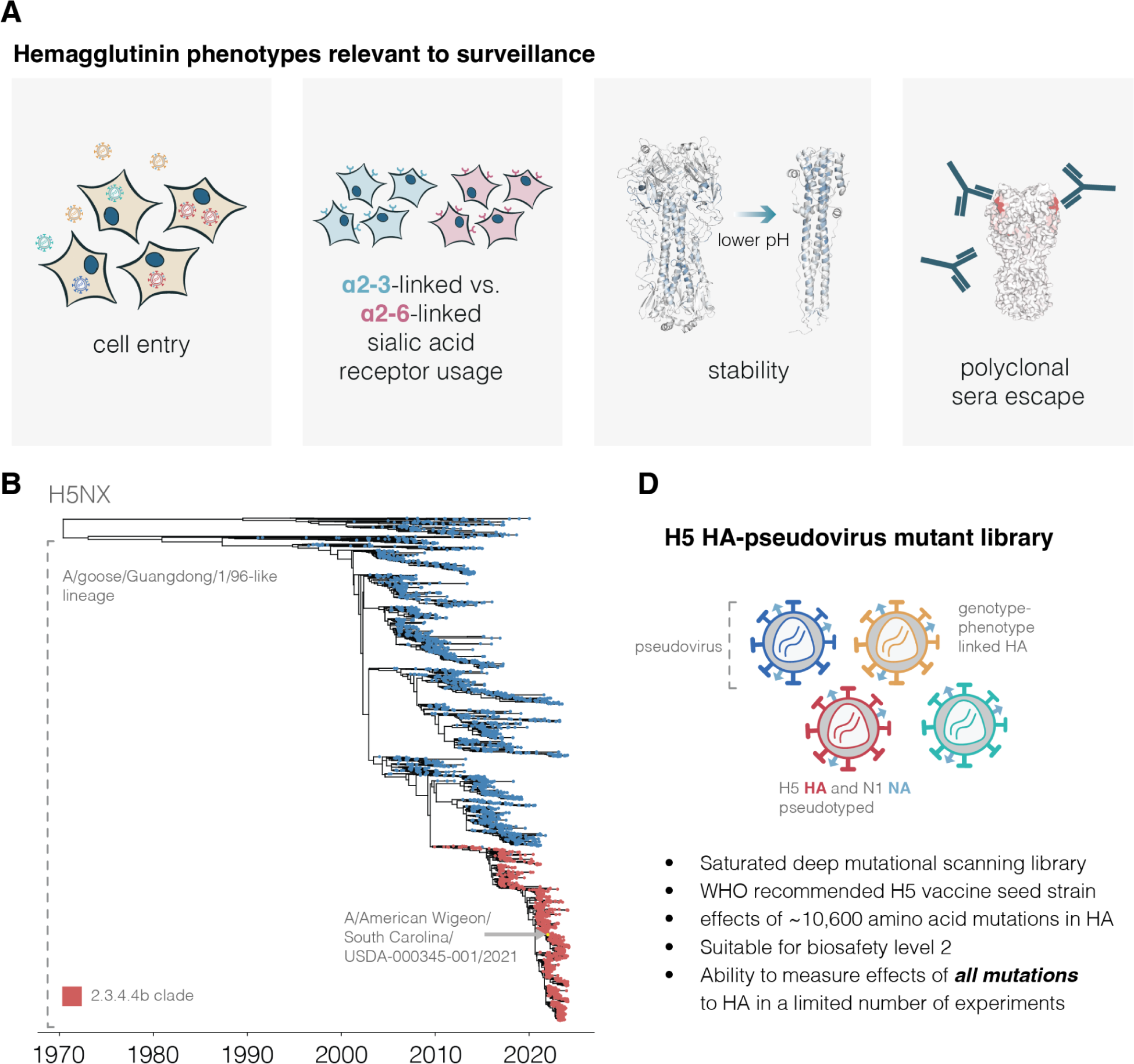
Deep mutational scanning of H5 HA. **A**, HA phenotypes relevant to pandemic risk that were measured in this study. **B**, Phylogenetic tree of H5Nx HAs. The A/goose/Guangdong/1/96-like lineage that was first identified as a highly pathogenic form of avian influenza in 1996 forms the large lower clade on the tree. The 2.3.4.4b clade that recently has spread globally is in red, and the A/American Wigeon/South Carolina/USDA-000345-001/2021 (H5N1) sequence used in our deep mutational scanning study is indicated. **C**, Schematic of a library of genotype-phenotype linked pseudoviruses. See **fig. S1** for more details on the library.

### H5 HA deep mutational scanning libraries

We designed H5 HA deep mutational scanning libraries to measure the effects of all 10,773 possible amino-acid mutations (567 sites ⨉ 19 amino-acid mutations/site). We created these libraries in the HA from the A/American Wigeon/South Carolina/USDA-000345-001/2021 (H5N1) strain, which is a WHO clade 2.3.4.4b candidate vaccine virus (*15*) (**Fig. 1B**). To generate pools of pseudoviruses with all the HA single amino-acid mutants, we used a lentivirus-based deep mutational scanning platform that generates pseudoviruses encoding different mutants of HA each linked to a nucleotide barcode in the lentiviral genome (*18*) (**fig. S1A-B**). These pseudoviruses also express the matched H5N1 neuraminidase (NA) on their surface; however, NA is expressed from a separate plasmid during pseudovirus production and only HA is mutagenized in our study (**fig. S1B**). The pseudoviruses can be studied at biosafety-level 2 since they encode no viral genes other than HA and so cannot undergo multi-cycle replication.

We produced duplicate pseudovirus libraries that contained 95% and 93% of the possible HA amino-acid mutations, respectively (**fig. S1C**). Most HAs contained a single amino-acid mutation, although a small fraction contained no or multiple mutations (**fig. S1D**).

### Effects of HA mutations on cell entry

To measure how mutations affect HA’s essential function of mediating viral entry into cells, we quantified HA-mediated pseudovirus entry into 293T cells (**fig. S2A**). These cells express both α2-3 and α2-6-linked sialic acids (*20*, *21*), which are typical receptors for avian and human influenza viruses, respectively. The measured effects of mutations on cell entry were highly correlated between replicate libraries (**fig. S2B**), demonstrating the reproducibility of the experiments.

There is wide variation in how HA mutations affect entry, with some sites tolerating many mutations and others under strong constraint (**Fig. 2** and interactive plot at https://dms-vep.org/Flu_H5_American-Wigeon_South-Carolina_2021-H5N1_DMS/cell_entry.html). Much of the constraint is shaped by how mutations affect HA’s folding and ability to mediate membrane fusion, since sites in the protein core including the stem helices in the fusion machinery (*22*) are especially intolerant to mutations (**Fig. 2B**). HA’s globular head domain is more mutationally tolerant than the stalk region, consistent with the head’s greater variability among natural influenza viruses and prior experimental work (*23–26*) (**Fig. 2B-C**). Additionally, the known antigenic regions of the head (*27*, *28*) are more tolerant to mutations than the rest of the head (**Fig. 2C**). However, there are some regions on HA’s surface that are intolerant of mutations (**Fig 2B**) and so represent attractive targets for efforts to design antibodies and other HA-directed therapeutics that are resistant to viral escape (*16*).

**Figure 2.**
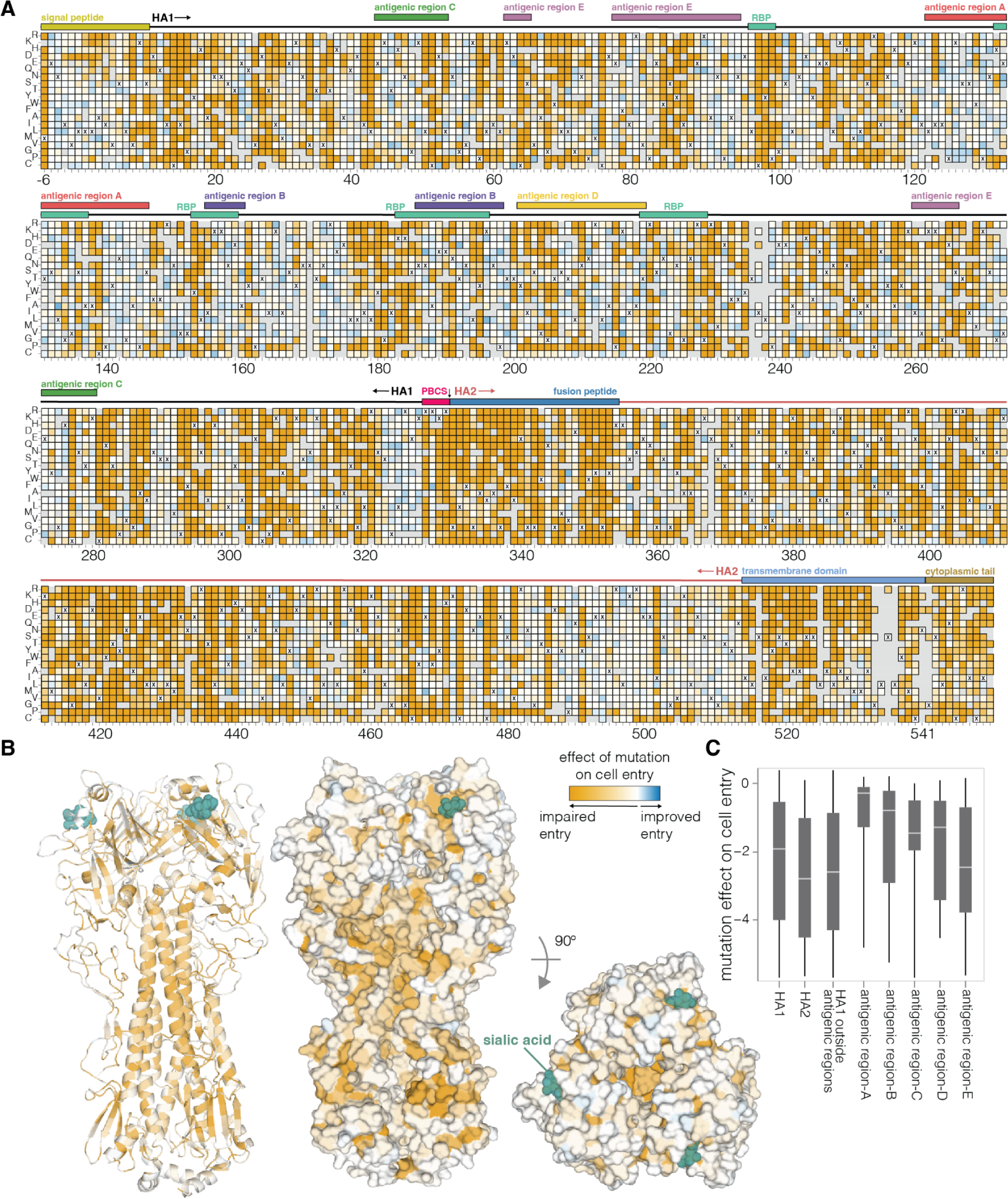
Effects of mutations on cell entry. **A**, Effects of mutations on pseudovirus entry into 293T cells. Orange indicates impaired entry, white parental-like entry, and blue improved entry. The x’s indicate the parental amino acid, and light gray indicates mutations that were not reliably measured in our experiments. See https://dms-vep.org/Flu_H5_American-Wigeon_South-Carolina_2021-H5N1_DMS/cell_entry.html for an interactive version of this heatmap that allows zooming and mousing over points for details. Sites are numbered using the H3 scheme (see https://dms-vep.org/Flu_H5_American-Wigeon_South-Carolina_2021-H5N1_DMS/numbering.html for details). **B**, HA structure (PDBs: 4KWM, 1JSO) colored by the average effect of amino-acid mutations at each site on cell entry (orange indicates sites where mutations tend to impair cell entry). **C**, Effects of mutations on cell entry for different domains and regions of HA. The boxes show the median and interquartile range, and more negative values indicate mutations adversely affect cell entry.

To validate that the effects of HA mutations measured in the pseudovirus deep mutational scanning reflect actual influenza virion cell entry, we generated conditionally replicative influenza viruses that lack the essential PB1 polymerase gene (*29*) and derive their other non-surface proteins from a lab-adapted strain (*30*), making them safe for study at biosafety-level 2 (**fig. S2C**). The effects of mutations on cell entry measured in the pseudovirus deep mutational scanning were highly correlated (R=0.9) with the titers of conditionally replicative influenza viruses carrying these same mutations (**fig. S2D**).

### Effects of HA mutations on α2-6-linked sialic acid receptor usage

Avian influenza viruses use α2-3-linked sialic acids as their preferred receptors, but human-transmissible strains typically prefer α2-6-linked sialic acids (*9–11*). To determine how mutations to HA affect its preference for these two receptors, we measured pseudovirus entry into 293 cells engineered to express primarily α2-3-linked or α2-6-linked sialic acids (hereafter referred to as α2-3 or α2-6 cells, see Methods for details on these cells) (*31*) by applying the workflow in **fig. S2A** separately to each cell line. While α2-6 cells still support limited infection by pseudoviruses with the unmutated H5 HA, titers on these cells are significantly lower than those on α2-3 cells because the parental H5 HA used in our deep mutational scanning is much more efficient at using α2-3-linked sialic acids (**fig. S3A**). Replicate measurements of mutational effects on entry made with the two different libraries were well correlated, again demonstrating the reproducibility of the measurements (**fig. S3B**).

Most HA mutations had similar effects on entry regardless of the cells used (**fig. S4**) since most mutations impact cell entry by affecting HA folding or membrane-fusion function (**Fig. 2B**), which are essential for entry into any cell. However, some mutations were more favorable for entry in α2-6 versus α2-3 cells (**Fig. 3**, **fig. S4**, interactive plots at https://dms-vep.org/Flu_H5_American-Wigeon_South-Carolina_2021-H5N1_DMS/a26_usage.html). The HA mutations that specifically enhanced entry into α2-6 versus α2-3 cells were mostly at a handful of sites in the sialic-acid binding pocket (**Fig. 3**). Multiple mutations to site Q226 strongly increased HA’s preference for α2-6-versus α2-3-linked sialic acids in our deep mutational scanning (**Fig. 3A**). These mutations include Q226L (throughout this manuscript we use mature H3 HA numbering, see https://dms-vep.org/Flu_H5_American-Wigeon_South-Carolina_2021-H5N1_DMS/numbering.html), which occurred in prior H3 and H2 HA pandemic viruses (*10*, *32*) and has previously been shown to enhance α2-6-linked sialic acid binding (*1*, *3*, *33*, *34*). Mutations at several other sites also markedly improved entry into α2-6 cells, including sites 137, 190, 193, 225, and 228 (**Fig. 3A-B**). Changes to some of these sites have occurred in previous pandemics (*10*, *32*, *35*, *36*) or been shown to enhance α2-6-linked sialic-acid binding (*33*, *34*, *37–39*) (**Fig. 3A-B**). Of the sites that strongly enhance entry into α2-6 cells, only a small number are accessible by single nucleotide change to the current 2.3.4.4b H5 HA gene sequence: these mutations include Q226L/R, E190Q/A, N224K/Y, G228A, and several others (**Fig. 3A**).

**Figure 3.**
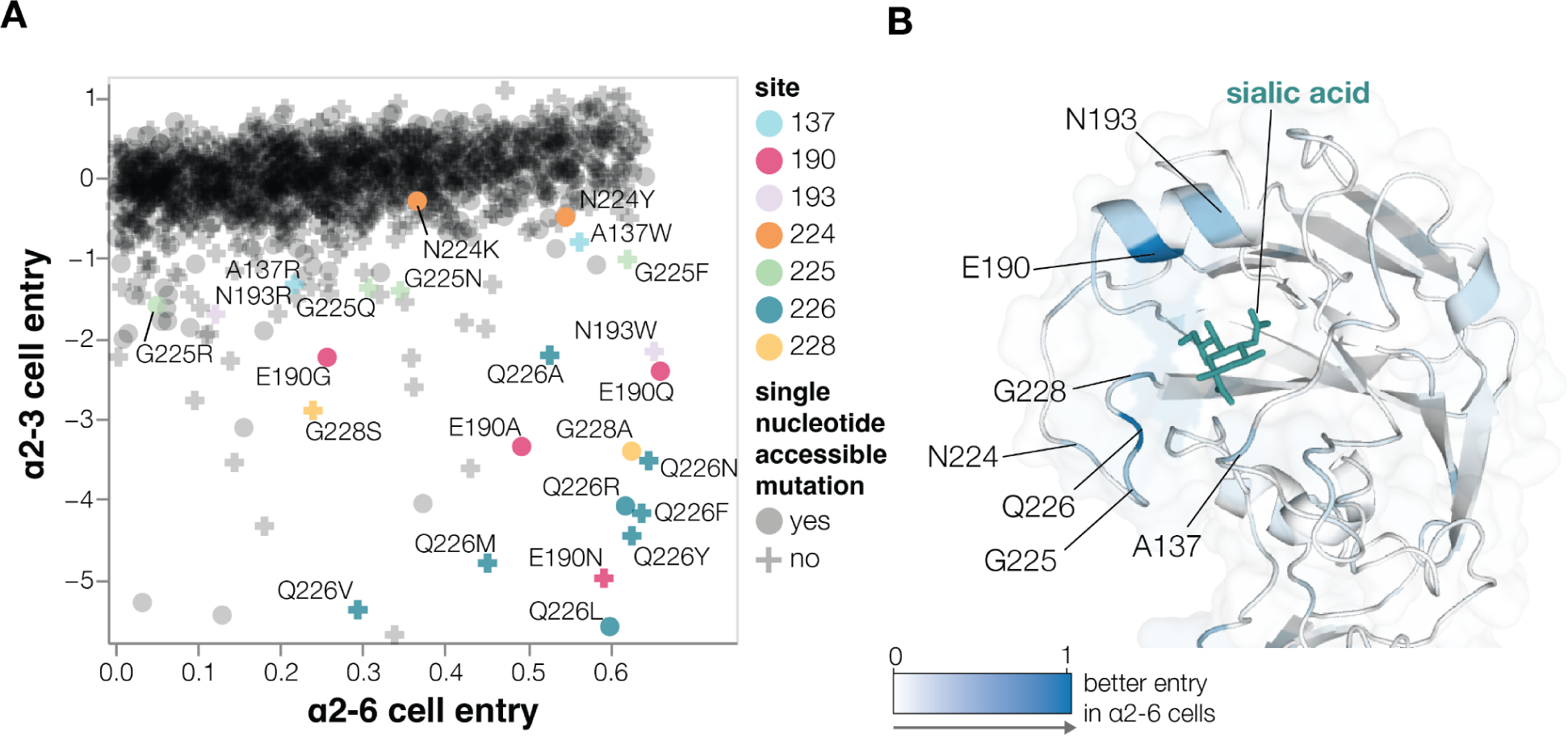
Effects of mutations on α2-6-linked sialic acid receptor usage. **A**, Effects of mutations on pseudovirus entry into 293 cells expressing α2-6-or α2-3-linked sialic acids. Each point is a different amino-acid mutation, with shapes indicating whether the mutation is accessible by a single-nucleotide change to the natural HA gene sequence. Sites where more than one mutation has large differences between α2-3 and α2-6 cell entry are labeled. Note that only mutations with a net favorable effect on entry into α2-6 cells are shown. **B**, Structure of the HA near the sialic-acid binding pocket (PDBs: 4KWM, 1JSO) with sites colored according to the average increase in α2-6 usage caused by mutations at that site. See https://dms-vep.org/Flu_H5_American-Wigeon_South-Carolina_2021-H5N1_DMS/a26_usage.html for interactive plots including a version of **A** that includes all mutations (not just those with net positive α2-6 cell entry) and an interactive heatmap of the change in α2-6 versus α2-3 usage for all mutations.

Note that while our experiments identify HA mutations that improve entry into the α2-6 293 cells, it is possible that these cells may not fully represent the abundance or diversity of different glycan structures in the human airways (*40–42*). In addition, in some cases multiple mutations may be needed to switch the preference of HA to α2-6 linked sialic acids (*43*).

### Effects of mutations on HA stability

Mutations that increase HA stability are associated with increased airborne transmissibility of influenza viruses (*1*, *3*, *44–49*). HA undergoes a similar irreversible conformational change in response to both acidic pH and elevated temperature (*22*), so the loss of viral infectivity at acidic pH is widely used as a measure of HA stability (*47*, *50*, *51*). To identify mutations that affect HA’s stability, we incubated the pseudovirus libraries in progressively more acidic pH buffers and quantified the retention of infectivity as a measure of HA stability (**Fig. 4A**).

**Figure 4.**
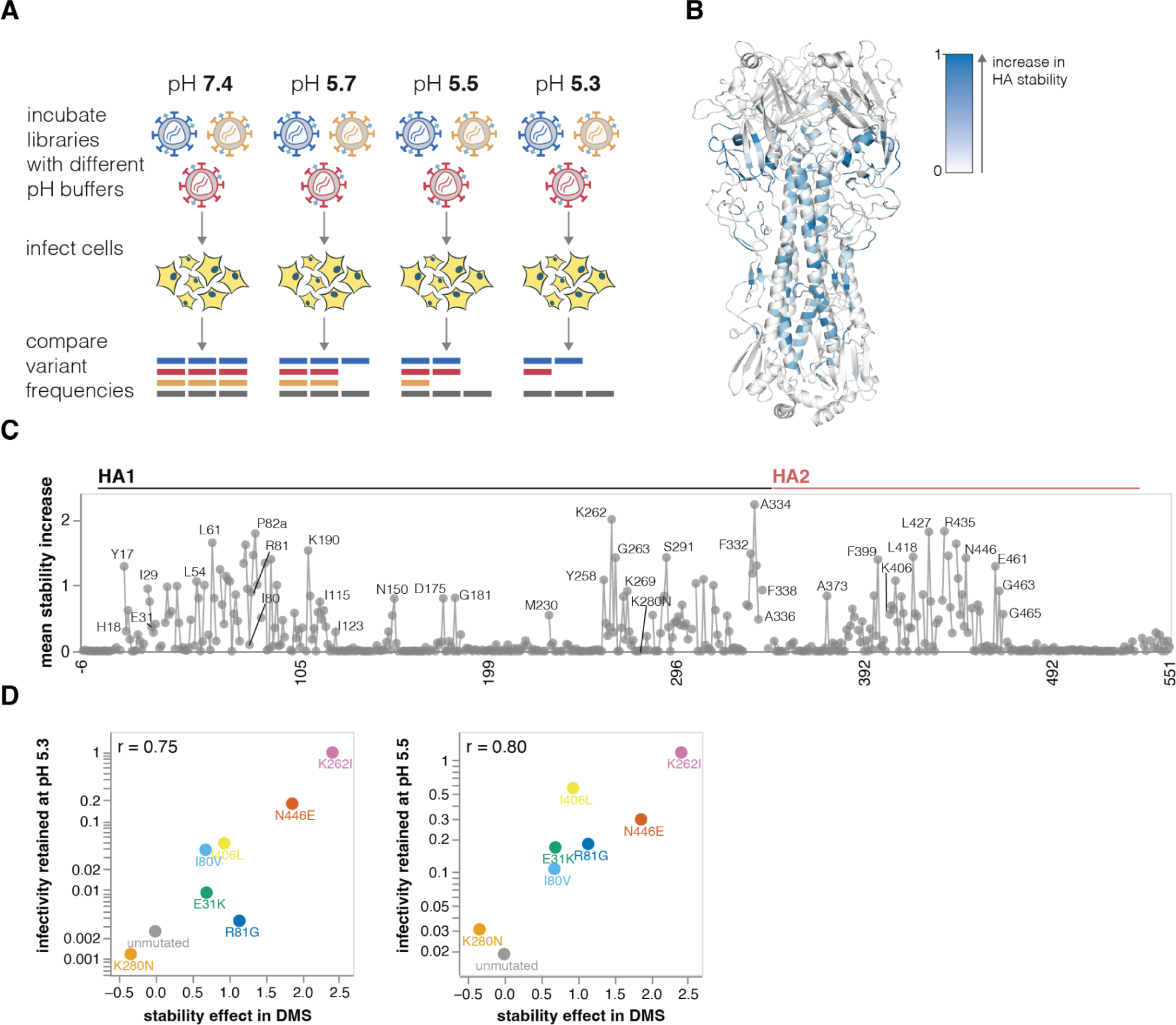
Effects of mutations on stability. **A**, To measure how mutations affect HA stability, the pseudovirus library was incubated in increasingly acidic buffers and then used to infect cells. Stability was quantified as the resistance of each mutant to inactivation by acidic pH. **B**, HA structure colored by the average increase in HA stability caused by mutations at each site, with blue indicating sites where mutations increase stability. **C**, The average increase in HA stability caused by mutations at each site. See https://dms-vep.org/Flu_H5_American-Wigeon_South-Carolina_2021-H5N1_DMS/stability.html for an interactive heatmap of the effects of all mutations that includes options to show destabilizing as well as stabilizing mutations. **D**, Validation of deep mutational scanning measurements by comparison to direct measurements of acid-induced inactivation of conditionally replicative influenza virions carrying the indicated mutations (see **fig. S5** for details). The mutations validated here were chosen to span a wide range of cell-entry effects.

Mutations at numerous sites increase HA stability (**Fig. 4B-C** and interactive plot at https://dms-vep.org/Flu_H5_American-Wigeon_South-Carolina_2021-H5N1_DMS/stability.html). Although these stabilizing mutations are widely distributed in primary sequence, they are concentrated in two regions of HA’s three-dimensional structure: the stem’s alpha helices and the base of the head domain (**Fig. 4B**). The spatial location of these mutations is consistent with the fact that acidic conditions or high temperature trigger an irreversible conformational change that involves rearrangement of the stem helices and displacement of the head domain (*22*, *52*). Overall, the deep mutational scanning both identified previously known stabilizing mutations (e.g., Y17H, A19T, H24Q, E31K, H110Y, and T318I (*1*, *2*, *14*, *46*, *53*)) as well as many additional mutations that increase stability (**Fig. 4C**).

We validated the deep mutational scanning stability measurements by generating conditionally replicative influenza virions (**fig. S2A**) carrying key mutations and quantifying their infectivity after acid treatment. The deep mutational scanning measurements correlated well with the stability of the influenza virions (**Fig. 4D, fig. S5**), validating the approach. We note that the map of stabilizing mutations could inform design of stabilized HA immunogens (*54*) for vaccines as well as inform viral surveillance.

### Effects of mutations on escape from serum neutralization

HA is the major target for neutralizing antibodies to influenza, but the protein can acquire mutations that escape neutralization by antibodies elicited by older strains (*55*). For potential pandemic influenza strains, the WHO designates candidate vaccine viruses (*15*), but these viruses can become poorly matched to newer strains as HA evolves. Assessing the impact of HA mutations on serum neutralization can identify when viruses have gained escape mutations that might merit updates to candidate vaccine viruses. The HA used for our mutant libraries is a WHO clade 2.3.4.4b H5 candidate vaccine virus (*15*). We used these libraries to measure how all HA mutations affect neutralization by polyclonal sera from mice or ferrets vaccinated or infected to elicit antibodies against clade 2.3.4.4b H5 HAs viruses (**fig. S6**).

The mutations that most strongly escaped serum neutralization were on the head of HA in previously defined antigenic regions (*56*) (**Fig. 5A-C** and interactive plots at https://dms-vep.org/Flu_H5_American-Wigeon_South-Carolina_2021-H5N1_DMS/escape.html). While the strongest escape mutations from both the mouse and ferret sera were in antigenic regions A and B at the top of the HA head, the specific sites where mutations caused the greatest escape did differ somewhat between the two species (**Fig. 5A-C**). There was also some animal-to-animal variation: for instance, most mice sera targeted antigenic regions A and B, but the serum of one mouse targeted antigenic region E lower on the HA head (**fig. S7**).

**Figure 5.**
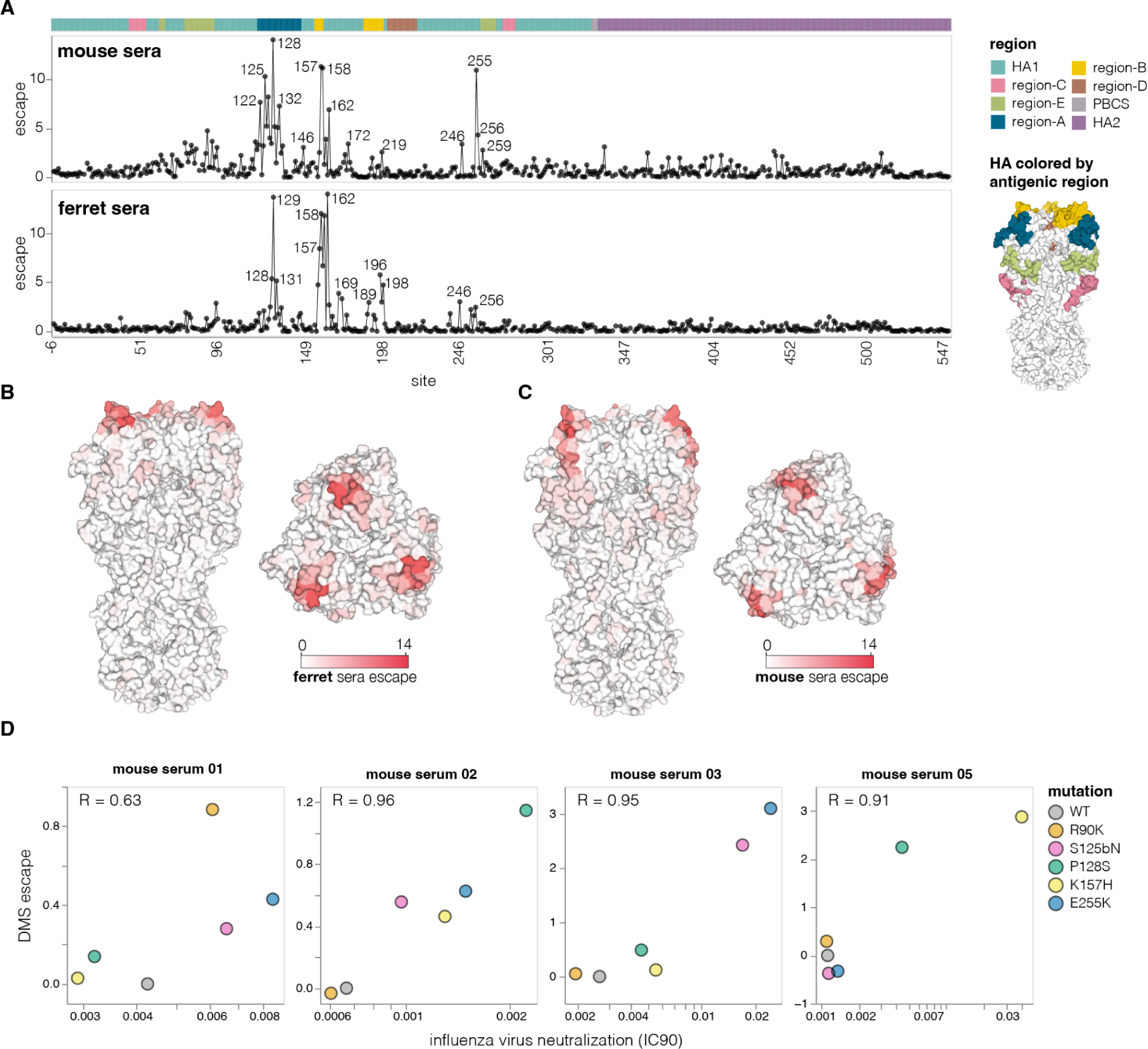
Effects of mutations on escape from serum neutralization. **A**, Total escape caused by all tolerated mutations at each HA site as measured in the deep mutational scanning, averaged across the sera of all animals from each species. The HA structure on the right shows the classically defined H3 antigenic regions (*56*). See **fig. S6B** for details on the number of animals and immunization methods. See https://dms-vep.org/Flu_H5_American-Wigeon_South-Carolina_2021-H5N1_DMS/escape.html for interactive plots showing escape for each animal. **B**, Average ferret sera escape overlaid on a surface representation of HA (PBD: 4KWM). **C**, same as in B but for mouse sera. **D**, Correlation between escape for different HA mutants measured with deep mutational scanning (DMS) versus the change in neutralization measured using conventional neutralization assays with conditionally replicative influenza virus.

To validate the escape mutations measured by deep mutational scanning, we performed standard neutralization assays using the conditionally replicative influenza viruses. The effects of mutations measured using deep mutational scanning correlated well with those measured using influenza-based neutralization assays (**Fig. 5D**). Therefore, even though pseudoviruses are generally more sensitive than actual influenza virions to neutralization (**fig. S8**), the mutation antigenic effects measured in the pseudovirus deep mutational scanning are recapitulated in actual influenza virions. Notably, several escape mutations that we validated (e.g., R90K, S125bN, P128S, E255K) are present among 2.3.4.4b clade sequences, illustrating the ongoing antigenic evolution of H5 HA.

### Phenotypic measurements inform sequence-based viral surveillance

Large numbers of H5 influenza sequences are collected during ongoing surveillance of outbreaks in various animals (*57–59*). One major goal of this surveillance is to identify viruses that acquire mutations that increase their risk to humans or alter their antigenicity in ways that necessitate updates to candidate vaccine viruses.

To use the large-scale phenotypic measurements described here to inform surveillance, we estimated the relevant HA phenotypes of clade 2.3.4.4b viruses as the simple sum of the phenotypic effects of their individual mutations. For instance, **Fig. 6A** shows a phylogenetic tree of recent viruses colored by their estimated escape from neutralization from sera from ferrets infected by 2.3.4.4b clade influenza virus; see https://nextstrain.org/groups/moncla-lab/h5nx/h5-dms/clade-2344b?c=ferret_sera_escape_dms_value for an interactive Nextstrain (*60*) version of the same tree. This tree enables immediate identification of subclades with antigenic mutations with reduced neutralization, including clades defined by mutations P128S, A160T/S and P162Q, which are present in some clusters of sequences from the ongoing dairy cattle outbreak in the US. These and other recent HA mutations also reduce neutralization by mouse sera (**fig. S9A**).

**Figure 6.**
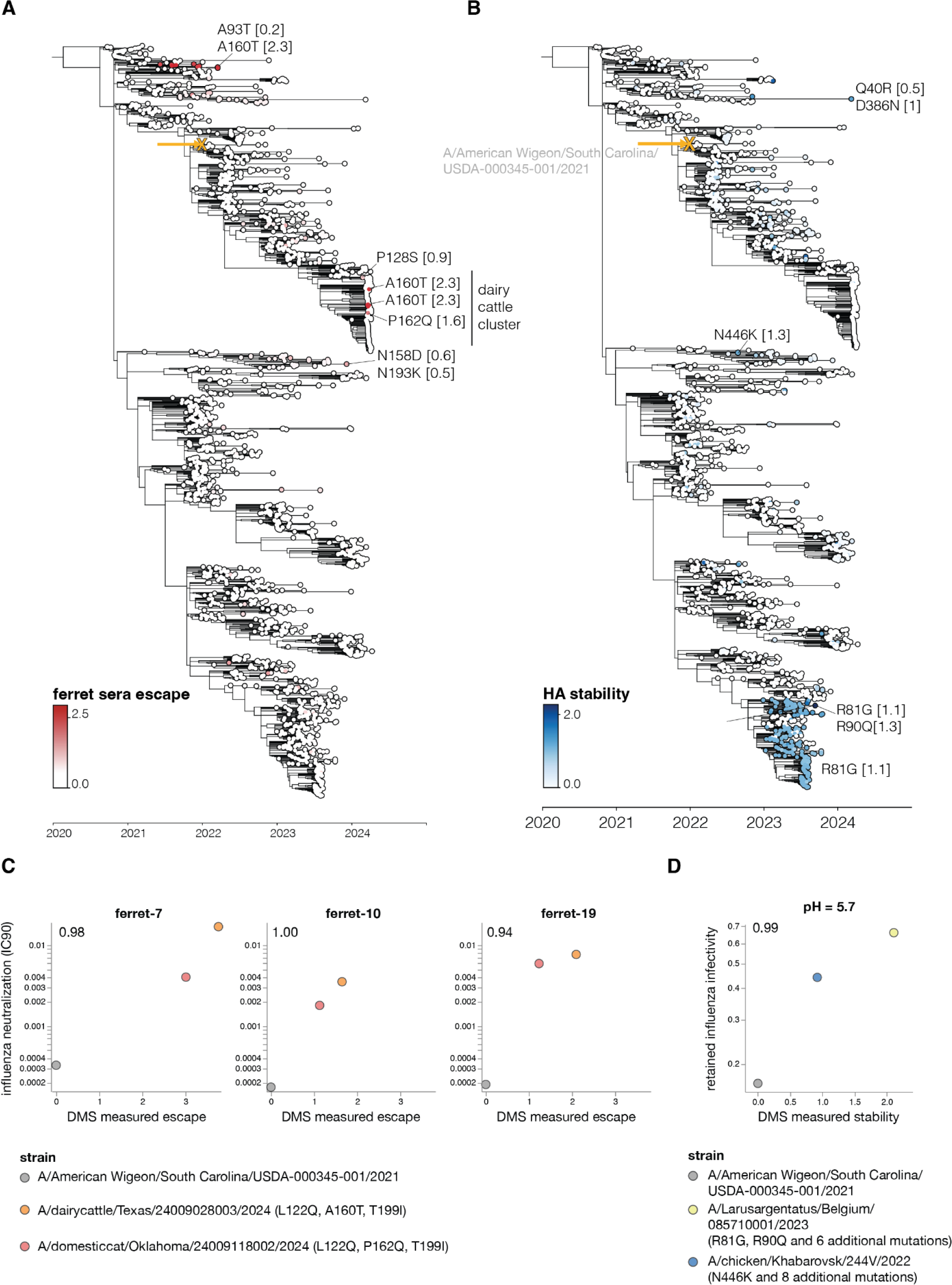
Estimated HA phenotypes of recent 2.3.4.4b strains. Phylogenetic trees of the HA gene of a subsample of H5 2.3.4.4b clade viruses collected from 2020 onwards. The trees are colored by (**A**) ferret serum escape and (**B**) HA stability. A phenotype is estimated for each sequence by summing the experimentally measured effects of all constituent HA mutations relative to the parental A/American Wigeon/South Carolina/USDA-000345-001/2021 strain used in the deep mutational scanning (indicated with a yellow “x” and arrow), only including mutations with positive effects in the sum. Some key mutations strongly affecting the displayed phenotypes are labeled on the tree, with the number in brackets indicating the effect of the mutation. See **fig. S9** for additional phenotypes mapped onto the same tree, and https://nextstrain.org/groups/moncla-lab/h5nx/h5-dms/clade-2344b for an interactive Nextstrain tree showing the phenotypes mapped onto various sets of sequences. **(C)** The HAs for strains carrying mutations that caused strong escape from ferret sera neutralization in the deep mutational scanning were assayed for neutralization by the ferret sera in conditionally replicative influenza virus neutralization assays **(fig. S10A)**. These scatter plots show the neutralization measured in the influenza virus assays (y-axis) versus the predicted escape calculated as the simple sum of the mutations in each HA relative to the strain used in the deep mutational scanning. **(D)** HAs from selected strains with stability-increasing mutations were assayed for their infectivity at different pHs using conditionally replicative influenza viruses. The scatterplot shows retained influenza virus infectivity at pH 5.7 relative to deep mutational scanning measured stability, which was calculated as a sum of stability effects for all mutations in a strain relative to the strain used in the deep mutational scanning. Retained infectivity at other pHs is shown in **fig. S10C**. The full list of HA amino-acid mutations in each strain is in **fig. S10D**

Similar estimates can be made for other HA phenotypes relevant to pandemic risk that were measured in our experiments. Some recent clade 2.3.4.4b HAs have acquired mutations that increase their stability, including groups of viruses sampled in 2023 with the R81G mutation (**Fig. 6B**). Likewise, a few (so far isolated) sequences on the tree have acquired mutations that enhance their usage of α2-6-linked sialic acids, including canine and human isolates from China that carry the Q226L mutation (**fig. S9B**).

To demonstrate the use of deep mutational scanning measurements for rapid identification of strains carrying potentially important mutations, we generated conditionally replicative influenza viruses with HAs from recent clade 2.3.4.4b strains that our experiments indicate have mutations that affect sera neutralization and HA stability (**Fig. 6, fig. S9, fig. S10**). Simply summing the effects of the individual mutations measured by pseudovirus deep mutational scanning accurately predicted the neutralization escape or stability of the actual influenza virions with HAs from different clade 2.3.4.4b strains (**Fig. 6C-D, fig. S9C, fig. S10A-C**), despite some of these HAs having up to 12 amino-acid mutations relative to the strain used in the deep mutational scanning (**fig. S10D**). The fact that some clusters of HA sequences from the recent dairy cattle outbreak carry mutations (P128S, A160T, or P162Q mutations) that appreciably reduce neutralization by mouse and ferret sera elicited by current candidate vaccine viruses underscores the importance of monitoring for potential antigenic drift relative to prototype vaccines.

Of course, sequence-based estimates of phenotypes from our deep mutational scanning are only approximate for viruses with more than one HA mutation relative to the parental strain used in our experiments, as epistasis can cause more diverged strains to have phenotypes different from the simple sum of the effects of their mutations. However, the deep mutational scanning accurately predicted serum neutralization and stability of multiple strains carrying as many as 12 amino-acid mutations relative to the strain used in our experiments (**Fig. 6C-D, fig. S9C, fig. S10**). This fact demonstrates that sequence-based phenotypic estimates informed by deep mutational scanning provide a method to rapidly assess newly observed viruses and prioritize them for more detailed experimental characterization, which remains important to validate findings and check for epistasis (*61*, *62*).

## Discussion

The last few decades have seen tremendous advances in viral sequencing. At the same time, experiments have advanced our understanding of molecular phenotypes of viral proteins including HA that contribute to pandemic risk (*1*, *3*, *9*, *63*). However, synthesizing these advances to make phenotypic assessments from sequence data remains challenging because only a small fraction of the mutations observed during viral surveillance have been experimentally characterized.

Here we take a step towards overcoming this challenge by experimentally measuring how nearly all mutations to H5 HA affect key molecular phenotypes. These measurements are valuable from both basic and applied standpoints. From a basic standpoint, they illuminate sequence-function relationships—for instance by comprehensively defining sites in HA’s stalk and head that contribute to its stability, and mutations that affect HA’s preference for α2-6-versus α2-3-linked sialic acids. From an applied standpoint, the measurements define which sites are constrained by the virus’s need to maintain HA’s essential cell entry function, thereby identifying targets for new approaches that design antibodies (*17*) and other therapeutics (*16*) to bind specific epitopes.

Importantly, our experiments are sufficiently comprehensive to inform phenotypic assessments of newly collected HA sequences. For example, it is known that HA’s stability and receptor specificity contribute to transmissibility (*1*, *3*, *44*), but actualizing this knowledge for surveillance has been hampered by an inability to assess how most HA mutations affect these phenotypes. By measuring the effect of nearly every HA mutation, our experiments enable immediate sequence-based estimation of stability and α2-6-linked sialic acid usage—thereby enabling rapid prioritization of viruses with HAs that merit further monitoring and experimental characterization (*61*, *62*).

Similarly, our measurements of how HA mutations affect serum antibody neutralization make it possible to immediately identify viruses that are mismatched to existing candidate vaccine viruses. For instance, the WHO recently released their antigenic measurements (*64*, *65*) on some HA mutations in clade 2.3.4.4b viruses associated with an ongoing dairy cattle outbreak in the United States (*64*). Their results are completely consistent with our deep mutational scanning: the two HA mutations basal to all cattle viruses have only modest antigenic effects in ferrets, but another mutation in a subset of viruses (A160T in H3 numbering, A156T in H5 numbering) causes an appreciable drop in neutralization. However, our data enable assessment of the antigenic effects of all the recently observed mutations, for instance showing that another mutation (P162Q in H3 numbering) found in a subset of cat viruses nested within the cattle viruses also causes substantial neutralization escape.

Our deep mutational scanning utilized pseudoviruses that encode only a single viral gene (HA), and so cannot undergo multi-cycle replication. Although we and others have previously performed deep mutational scanning using replicative influenza viruses with genes from biosafety-level 2 strains (*24*, *25*, *66*), we abandoned that approach in favor of pseudoviruses to avoid biosafety concerns associated with mutating replicative viruses with HA from a potential pandemic strain (*67*). We acknowledge that the use of pseudoviruses does not eliminate information hazards (*68*, *69*); however, specific examples of mutations that promote airborne-transmissibility of H5N1 by increasing HA stability and α2-6-linked sialic-acid usage have already been publicly described (*1*, *3*), so safely measuring how other HA mutations affect these phenotypes benefits surveillance efforts without creating qualitatively new information risks.

There are several caveats when applying the measurements reported here to viral surveillance. First, HA is just one of several influenza proteins (*4*, *6–8*) where mutations affect pandemic risk. Second, the pseudoviruses we used do not fully capture the balance between HA and NA protein activities that contributes to influenza virus fitness (*70–72*). Third, the cell lines used in our experiments probably do not fully recapitulate the distribution of glycan receptors in the human airways. Fourth, we measured how mutations affected neutralization by mice and ferret sera, which may have different specificities than human sera (although our experiments could be easily applied to human sera if it becomes available). Finally, we measured the effects of individual mutations, but mutations may sometimes combine in non-additive ways, particularly in HAs that are substantially diverged from the parental HA used in our experiments (*73–76*). Nonetheless, the experimental results reported here demonstrate that the pseudovirus deep mutational scanning can accurately predict the neutralization and stability of other clade 2.3.4.4b HAs.

Overall, our study provides the first comprehensive measurements of how HA mutations affect key molecular phenotypes that contribute to the pandemic risk posed by H5 influenza. Combining these measurements with rapid sequencing of ongoing H5 influenza outbreaks can improve monitoring of ongoing viral evolution, as well as inform the choice of candidate vaccine viruses, the targeting of designed therapeutics (*16*, *17*), and the prioritization of viral strains for additional experimental study (*61*, *62*).

## Acknowledgments

We thank Henrik Clausen and Yoshiki Narimatsu from Copenhagen Center for Glycomics for sharing the α2-3 and α2-6 cells used in this study. We thank Charles Russell for helpful advice about HA stability. We thank Brendan Larsen for help with the homepage hosting the interactive plots. We thank all the scientists who contributed H5 HA sequences shown in the phylogenetic trees in this paper to Genbank or GISAID.

## Funding

NIH/NIAID grant R01AI141707 (J.D.B.)

NIH/NIAID grant R01AI165821 (J.D.B.)

NIH/NIAID grant AI47029 (L.H.M.)

NIH/NIAID contract 75N93021C00015 (S.E.H., J.D.B., and L.H.M.)

Gift from Open Philanthropy (to N.P.K.)

UK Medical Research Council/Department for Environment (J.Y., Y.L., T.P.P) Food and Rural Affairs (Defra, UK) (J.Y., Y.L., T.P.P)

FluTrailMap-One Health consortium grant MR/Y03368X/1 (J.Y., Y.L., T.P.P)

Biotechnology and Biological Sciences Research Council (BBSRC)/DEFRA ‘FluTrailMap’ consortium grant BB/Y007298/1 (J.Y., Y.L., T.P.P)

BBSRC via the Pirbright Institute’s Strategic Programme Grant BBS/E/PI/230002A (T.P.P) NIH grant S10-OD-020069 (Fred Hutch Scientific Computing)

NIH grant S10-OD-028685 (Fred Hutch Scientific Computing).

Howard Hughes Medical Institute (J.D.B.)

## Author contributions

Conceptualization: B.D., J.D.B., T.P.P., L.H.M.

Methodology, B.D., J.D.B., Y.L., L.H.M.

Experiments: B.D., J.J.A., C.F. J.Y., A.D.

Computational analysis: B.D., J.D.B, W.W.H., J.T.O., L.H.M.

Writing – original draft, B.D. and J.D.B.

Writing – review & editing, all authors

Resources, A.L.V.B., R.W., N.P.K., S.E.H.

Supervision J.D.B.

Funding acquisition, J.D.B.

## Data and materials availability

All data generated in this paper can be viewed interactively at https://dms-vep.org/Flu_H5_American-Wigeon_South-Carolina_2021-H5N1_DMS. All analyses can be reproduced using the pipeline available at https://github.com/dms-vep/Flu_H5_American-Wigeon_South-Carolina_2021-H5N1_DMS. See Materials and Methods for the full description of the computational pipeline used for analysis. Raw sequencing files are available under BioProject PRJNA1123200.

## Material and Methods

### Data availability and computer code

The data reported in this paper are available in both interactive plots and spreadsheets of numerical values. See https://dms-vep.org/Flu_H5_American-Wigeon_South-Carolina_2021-H5N1_DMS for a webpage that hosts interactive plots that allow easy visualization and querying of the data. We recommend readers initially examine the summary page (https://dms-vep.org/Flu_H5_American-Wigeon_South-Carolina_2021-H5N1_DMS/summary.html), and then query the additional more detailed plots provided for individual phenotypes as needed. Note that the plots provided for individual phenotypes on the homepage show the measurement for each individual replicate library when you mouse over the heatmaps, which can provide a useful way to assess the reproducibility associated with any specific measurement. See https://github.com/dms-vep/Flu_H5_American-Wigeon_South-Carolina_2021-H5N1_DMS/blob/main/results/summaries/phenotypes.csv for the numerical values of the measurements with our recommended filtering for only high-confidence measurements; the main interactive homepage mentioned above also contains links to additional CSV files with less filtered values and per-replicate measurements (although be sure you understand the filters before using those).

The data can be viewed interactively in the context of the H5 HA protein structure using *dms-viz* (*77*) at https://dms-viz.github.io/v0/?data=https%3A%2F%2Fraw.githubusercontent.com%2Fdms-vep%2FFlu_H5_American-Wigeon_South-Carolina_2021-H5N1_DMS%2Fmain%2Fresults%2Fdms-viz%2Fdms-viz.json

See https://github.com/dms-vep/Flu_H5_American-Wigeon_South-Carolina_2021-H5N1_DMS for a GitHub repository that contains all of the computer code used in the analysis. The repo provides instructions on how to run the pipeline either by starting with raw fastq files, which can be downloaded from BioProject PRJNA1123200, or from already pre-build barcode count files. Most of the analysis was performed using dms-vep-pipeline-3 (https://github.com/dms-vep/dms-vep-pipeline-3), version 3.13.0. Note that rerunning the pipeline on different computers may give slightly different numbers due to platform-specific differences in floating point precision.

See https://nextstrain.org/groups/moncla-lab/h5nx/h5-dms/usa-clade-2344b for interactive Nextstrain (*60*) trees with the deep mutational phenotypes used to color the nodes; you can use the dropdown boxes at left to select which phenotype to use to color the nodes (the *Color By* box) or what set of sequences to show (eg, *all-clades*, *clade-2344b*, or *usa-clade-2344b*). The parental sequence used for the deep mutational scanning is marked with an “x”, and the estimated phenotypes of other sequences are expected to become less accurate the more diverged those sequence are from this parental sequence. Code for annotating DMS data onto phylogenies is available at https://github.com/moncla-lab/annotate-dms.

### HA sequence numbering

Throughout we use mature H3 numbering of the HA sequence. See https://dms-vep.org/Flu_H5_American-Wigeon_South-Carolina_2021-H5N1_DMS/numbering.html for details and for a file that interconverts various numbering schemes.

### Design of deep mutational scanning libraries

Deep mutational scanning libraries were designed in the background of A/American Wigeon/South Carolina/USDA-000345-001/2021 strain. The HA gene was codon optimized for use in the lentiviral backbone; see https://github.com/dms-vep/Flu_H5_American-Wigeon_South-Carolina_2021-H5N1_DMS/blob/main/library_design/pH2rU3_ForInd_H5_H5N1_American_Wigeon_genscript_T7_CMV_ZsGT2APurR.gb for the backbone plasmid containing the HA gene. The libraries were designed to have every possible amino acid mutation to HA and every other STOP codon in the first 20 positions of HA. Single mutant libraries were ordered from Twist Biosciences. Final quality control data for libraries produced by Twist is at https://github.com/dms-vep/Flu_H5_American-Wigeon_South-Carolina_2021-H5N1_DMS/blob/main/library_design/Q-248900_Final_QC_Report_for_twist_library.xlsx.

### Deep mutational scanning library plasmid library production

18 out of 567 codon sites in the library failed during the variant library production by Twist Biosciences. 8 of these sites were in the transmembrane domain and 10 were in the HA ectodomain. We decided to mutagenize the 10 failed ectodomain sites in-house and add them to the library to ensure complete mutagenesis of the ectodomain; we did not attempt to re-add the failed sites in the transmembrane domain since mutations in that region are unlikely to affect the HA phenotypes of interest for this study. To add mutations at the missing ectodomain sites, forward and reverse primers with NNG/NNC codons were designed and ordered from IDT, sequences for these primers can be found at https://github.com/dms-vep/Flu_H5_American-Wigeon_South-Carolina_2021-H5N1_DMS/blob/main/library_design/mutagenesis_primers.csv. Forward and reverse primer pools were made by pooling primers at equimolar ratios to a final concentration of 5 µM. Linear HA template was produced by digesting the lentivirus backbone plasmid encoding HA (see above for plasmid map) with NotI and XbaI enzymes and gel purifying the digestion reaction. A PCR mutagenesis reaction was then performed as described previously (*18*) with a few changes, Namely, 12 PCR cycles were used for the mutagenesis reaction and after the joining PCR reaction was purified, an additional Dpn digest was performed for 20 min in order to remove any potential wild-type template. Mutagenised HA fragments were then pooled together with the linear library pool produced by Twist Biosciences at a molar ratio proportional to the number of mutants in each pool so that each codon variant would have an equal representation in the final library. The pooled library was then split into two reactions (creating library-1 and library-2, which were then handled independently through the rest of the deep mutational scanning) and barcoded with primers containing a random 16-nucleotide sequence downstream of the stop codon as described previously (*18*). To produce plasmid libraries, mCherry-containing lentivirus backbone (plasmid map at https://github.com/dms-vep/Flu_H5_American-Wigeon_South-Carolina_2021-H5N1_DMS/blob/main/library_design/3137_pH2rU3_ForInd_mCherry_CMV_ZsGT2APurR.gb) was digested with XbaI and MluI restriction enzymes and barcoded linear HA libraries were inserted into the lentivirus backbone using NEB HiFi assembly kit (NEB E5520S). Assembled plasmid libraries were then electroporated into electrocompetent 10b cells (NEB C3020K) and incubated overnight on ampicillin containing LB plates. Ten electroporation reactions were performed per library, producing > 3.7 M CFUs per library in total. High colony numbers are required to have a large barcode diversity in the plasmids in order to prevent the same barcode becoming associated with multiple different HA variants during lentiviral recombination during subsequent steps in the production of the pseudovirus libraries. The colonies were scraped from the LB plates and plasmid library pools were purified using a QIAGEN Plasmid Maxi Kit (Cat. no. 12162).

### Production of cell-stored deep mutational scanning libraries

Cells storing the deep mutational scanning libraries as a single integrated HA-containing proviral genome per cell (**fig. S1A-B**) were produced as described previously (*18*) with a few changes. In brief, VSV-G pseudotyped viruses were produced by transfecting two 6-well plates of 293T cells with the lentiviral backbone containing the barcoded HA deep mutational scanning libraries (1 ug per well), lentiviral Gag/Pol, Tat and Rev helper plasmids (BEI Resources: NR-52517, NR-52519, NR-52518, 0.25 ng per well each), a VSV-G expression plasmid (0.25 ng per well, plasmid map at https://github.com/dms-vep/Flu_H5_American-Wigeon_South-Carolina_2021-H5N1_DMS/blob/main/library_design/29_HDM_VSV_G.gb), and plasmid expressing the neuraminidase (NA) gene (1 µg per well, plasmid map at https://github.com/dms-vep/Flu_H5_American-Wigeon_South-Carolina_2021-H5N1_DMS/blob/main/library_design/3405_HDM_N1_native_H5N1_turkey_Indiana.gb). Neuraminidase gene came from A/turkey/Indiana/22-003707-003/2022 strain, this strain also belongs to 2.3.4.4b clade and has the same HA sequence but differs in NA sequence by 2 amino acids relative to the NA for the strain used in this study. This NA was used because it provided high pseudovirus titers and NA from the A/American Wigeon/South Carolina/USDA-000345-001/2021 strain was never tested. Note that while VSV-G pseudoviruses do not require NA for virus release or entry, the produced virions will also have HA expressed from the lentiviral backbone on their surface, so NA expression is needed to prevent HA from binding the virions to the producer cell surface. This is particularly important for a mutant library, as different HA mutants will have different expression and sialic-acid binding properties, which could lead to a different tendency of different virions to be retained on the cell surface by HA if NA is not added at a high excess. The VSV-G pseudotyped viruses were then used to infect 293T-rtTA cells (using a specific clone we have previously found to yield good pseudovirus titers (*18*)) at an infection rate of 0.4% so that most transduced cells receive only a single integrated proviral genome. Transduced cells were then selected using puromycin leading to a population of cells in which each cell contains an integrated proviral genome encoding a single barcoded variant of HA (**fig. S1A-B**). These cells were then frozen in ∼20 M cell aliquots and stored long term in liquid nitrogen for further use.

### Long-read PacBio sequencing for variant-barcode linkage

To link barcode sequences with mutations in HA (**fig. S1A**). We performed long-read PacBio sequencing of lentiviral genomes in virions produced from cell-stored libraries as described previously (*18*). To do so 90 million cell-stored library cells were plated in a 3-layer flask and next day transfected with lentiviral helper plasmids (26.4 µg each), NA expression plasmid (10.8 µg) and VSV-G expression plasmid (11.25 µg) to produce VSV-G pseudotyped virions. At 30 hours after transfection, the virus-containing cell supernatant was collected and concentrated 15-fold using a protein column concentrator (Pierce™ Protein Concentrator Cat. no. 88537). Around 15 million transcription units of the concentrated virus stock were then used to infect 293T cells, and non-integrated reverse-transcribed lentiviral genomes were recovered by miniprepping the 293T cells as described previously (*18*). Eluted genomes were then split into two PCR reactions each of which added a unique tag for detecting strand exchange, which would prevent correct mutation-barcode linkage. Aliquots from the two PCR reactions were then pooled together and another round of PCR was performed. In each PCR round we minimized the number of reaction cycles in order to limit strand exchange. Long read sequencing was then performed on PacBio Sequel IIe machine. The PacBio circular consensus sequences (CCSs) were then aligned to the target amplicon (https://github.com/dms-vep/Flu_H5_American-Wigeon_South-Carolina_2021-H5N1_DMS/blob/main/data/PacBio_amplicon.gb) using *alignparse* (*78*), and consensus sequences for the HA variant associated with each barcode were determined requiring at least two CCSs per barcode. The final barcode-variant lookup table can be found at https://github.com/dms-vep/Flu_H5_American-Wigeon_South-Carolina_2021-H5N1_DMS/blob/main/results/variants/codon_variants.csv. Library 1 had 40,458 variants and library 2 had 34,495 variants, with an average of 1.1 mutations per HA (**fig. S1C-D**).

### Mutation effects on cell entry

Cell entry effects of each mutation in the library were determined as described previously (*18*) with some modifications (see also **fig. S2A**). HA pseudotyped library viruses were produced from 5-layer flasks in the presence of doxycycline (which induces HA expression from lentiviral backbone) and using 44 µg of each lentiviral plasmid and 18 µg of NA expression plasmid. Virus-containing supernatant was recovered 48 hours post transfection and concentrated 7.5-fold using either protein column concentrator or Lenti-X™ concentrator (TakaraBio, Cat. no. 631232) according to manufacturer’s instructions. To measure effects of mutations on cell entry, 293T cells were infected with either libraries pseudotyped with VSV-G envelope (produced as described in the previous methods subsection) or HA pseudotyped libraries. For HA pseudotyped library infections 293T cells were plated in the presence of 2.5 µg/ml of amphotericin B a day before, which increases library virus titer by ∼4 fold, and infections were performed in the presence of 1 µM oseltamivir in order to inhibit NA and prevent cleavage of receptors on target cell surface. For VSV-G pseudotyped library infections, ∼15 million transcription units were used and for H5 pseudotyped library infections ∼5 million transcription units were used. At 12-15 hours post infection cells were collected, nonintegrated lentiviral genomes were recovered using QIAprep Spin Miniprep Kit (Qiagen, cat. no. 27104) and amplicon libraries for barcode sequencing were prepared as described previously (*8*).

Cell entry scores for each variant were calculated using log enrichment ratio: 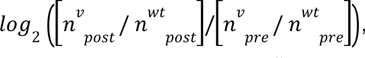, where *n^v^_post_* is the count of variant *v* in the H5-pseudotyped infection (post-selection condition), *n^v^_pre_* is the count of variant *v* in the VSV-G-pseudotyped infection (pre-selection condition), and *n^wt^_post_* and *n^wt^_post_* are the counts for wild-type variants. Positive cell entry scores indicate that a variant is better at entering the cells compared to the unmutated parental HA, and negative scores indicate entry worse than the parental HA.

To calculate the mutation-level cell entry effects, a sigmoid global-epistasis (*79*) function was fitted to variant entry scores after truncating the values at a lower bound of −6, using the *multi-dms* software package (*80*). The rationale for fitting the global epistasis models is that some of the variants have multiple HA mutations, and so the global epistasis model enables deconvolution of the mutation effects from these multiply mutated variants as well as just the single mutants. An interactive heatmap showing the effect of each mutation on cell entry is at https://dms-vep.org/Flu_H5_American-Wigeon_South-Carolina_2021-H5N1_DMS/cell_entry.html. For the mutation effects, values of zero mean no effect on cell entry, negative values mean impaired cell entry, and positive values mean improved cell entry. For the plots in this paper, we only show by default mutations observed in average of at least two barcoded variants per library and measured in both of the duplicate libraries.

### Effects of mutations on α2-6-versus α2-3-linked sialic acid cell entry

To identify mutations that increase α2-6-linked sialic-acid receptor preference we performed cell entry selections like the ones described above for 293T cells using cells that express either α2-3-linked or α2-6-linked sialic acids (see the Cell lines methods subsection below for the full description of these cell lines). For cell entry measurements on α2-3 cells, ∼5 million transcription units of HA pseudotyped library virus were used, while for α2-6 cells ∼20 million transcription units of the library pseudovirus were used, with the titers (transcriptional units) determined on each cell line separately. The reason higher titers could be used for the α2-3 cells is because virus titers on α2-6 cells are significantly lower (**fig. S3A**). Cell entry scores for each mutation in the library were calculated the same way as for 293T cells except that VSV-G pseudotyped virus infection on α2-6 cells served as pre-selection condition for α2-3 cell entry, and VSV-G infection on α2-3 cells served as pre-selection condition for α2-6 cell entry. The reason for this change in pre-selection condition is because H5 HA pseudotypes just as effectively as VSV-G and so viruses pseudotyped with VSV-G will have H5 HA on their surface which will be expressed from lentiviral backbone due the the presence of Tat during pseudovirus production. Because of this, variants of H5 HA with mutations that enhance α2-3 or α2-6-linked sialic acid usage would also lead to VSV-G pseudotyped virus enter those respective cells better which would in turn lead to lower cell entry scores for such variants. Using unmatched pre- and post-selection conditions should mitigate this entry bias, although in practice while entry scores for key sialic acid preference determining mutations do change when pre-selection and post-selection conditions are not matched, the differences are fairly small. To calculate the increase in α2-6-linked sialic acid usage we subtracted α2-3 cell entry mutation effects from α2-6 cell entry mutation effects, such that positive differences indicate better entry into the α2-6 cells. Interactive plots showing ell entry scores for α2-3 cells are at https://dms-vep.org/Flu_H5_American-Wigeon_South-Carolina_2021-H5N1_DMS/htmls/293_SA23_entry_func_effects.html, cell entry scores for α2-6 cells are at https://dms-vep.org/Flu_H5_American-Wigeon_South-Carolina_2021-H5N1_DMS/htmls/293_SA26_entry_func_effects.html and differences between α2-3 and α2-6 cell entry are interactivelydisplayed at https://dms-vep.org/Flu_H5_American-Wigeon_South-Carolina_2021-H5N1_DMS/a26_usage.html.

### Effects of mutations on HA stability

To measure the effects of mutations on HA stability, we followed the workflow outlined in (**Fig. 4A**). Specifically, ∼5 million transcription units of the pseudovirus library were incubated with D10 media or citrate-based acidic buffers. Buffers were prepared at 0.1 M by mixing together sodium citrate dihydrate and citric acid solutions and adjusting to pHs 5.7, 5.5, or 5.3 using hydrochloric acid or sodium hydroxide. Pseudovirus libraries were incubated for 1 hour at 37 °C at 1:12 library to buffer volume ratio. After incubation, libraries were then concentrated using 100k Amicon spin columns (Milipore, UFC910008) by spinning for 15 min at 1500 g to ∼1 ml volume and resuspended in 11 ml of PBS to exchange and neutralize acidic buffers. Viruses were spun down again for 20 minutes at 1500 g to reduce sample volume and exchange PBS for 2 ml of D10 media. Oseltamivir was added to the libraries at 1 µM and they were then used to infect 293T cells that were plated a day before in the presence of 2.5 µg/ml of amphotericin B. At 12-15 hours post infection unintegrated, lentiviral genomes were recovered from cells using Qiagen Miniprep Kit. During the miniprep lysis step (P2 buffer) 0.003 ng of DNA standard was added to a final molar concentration of ∼1% of recovered lentiviral DNA (this was calculated based on the number of expected non-integrated lentiviral genomes recovered given the amount of virus used for infections). The DNA standard consisted of a plasmid encoding lentiviral backbone with a barcoded mCherry gene (plasmid map for non neutralizable standard is at https://github.com/dms-vep/Flu_H5_American-Wigeon_South-Carolina_2021-H5N1_DMS/blob/main/library_design/3068_ForInd_mC_BCs_pool1.gb and barcodes in this standards can be found at https://github.com/dms-vep/Flu_H5_American-Wigeon_South-Carolina_2021-H5N1_DMS/blob/main/data/neutralization_standard_barcodes.csv), and the purpose of this standard is to enable conversion of relative sequencing counts to absolute amount of each barcoded pseudovirus variant (relative to the standard) at each pH condition. Note that unlike for sera selections (described below) where we use VSV-G or RdPro pseudotyped viruses as non-neutralizable standard, here we choose to use DNA-based standard because VSV-G or RdPro may be affected by the pH treatment. Lentiviral genomes were then used to make amplicon libraries for Illumina sequencing as described previously (*18*).

The sequencing counts were converted to the fraction infectivity retained at each pH condition relative to D10 by using the counts of each variant relative to the DNA standard at each condition. The *polyclonal* (*81*) software (v6.11) was used to analyze mutation-level retained infectivity at each pH and to fit a pH-dependent neutralization curve while estimating the effect of each mutation on stability. An example notebook of this analysis can be found at https://dms-vep.org/Flu_H5_American-Wigeon_South-Carolina_2021-H5N1_DMS/notebooks/fit_escape_stability_Lib2-231002-pH.html.

### Effects of mutations on serum neutralization

To measure the effects of mutations on neutralization by polyclonal mouse or ferret sera, we followed the workflow in **fig. S6A**, which parallels previously described work (*18*, *82*). Specifically, ∼2.5 million transcription units of the library virus were mixed with non-neutralizable standard so that the final amount of DNA from the non-neutralized standard would be ∼ 1% in the absence of serum selection. For mouse sera VSV-G pseudotyped viruses were used as non-neutralizing standards, while for ferret sera RdPro pseudotyped viruses were used. Both of these non-neutralizable standards and their generation have been described previously (*82*). The reason different neutralizing standards were used for the two types of sera is because ferret sera non-specifically neutralizes VSV-G virus but does not neutralize RdPro virus, while mouse sera does not neutralize either; as VSV-G pseudovirus is easier to produce we opted to use it for mice sera instead of the RdPro. The choice of the non-neutralizable standard should not influence sera selection results as both pseudoviruses carry identical lentiviral genomes. At the infection stage, 1 µM of oseltamivir was also added. Library viruses were then incubated with three sera dilutions that were measured to reach IC99, IC99*4 and IC99*16 neutralization using luciferase-based pseudovirus neutralization assay for 45 min at 37 °C. Strongly neutralizing dilutions of sera need to be used in these assays in order to select for only the variants that can escape polyclonal sera due to mutations they carry, and multiple sera dilutions are used to cover a greater dynamic range. After incubation with serum, 293T cells that were plated in the presence of 2.5 µg/ml of amphotericin B a day before were infected with library viruses. At 12-15 hours post infection cells were collected, nonintegrated lentiviral genomes were recovered using QIAprep Spin Miniprep Kit and amplicon libraries for barcode sequencing were prepared as described previously (*18*).

The data were analyzed as described previously (*18*, *82*). Briefly, the fraction infectivity retained at each serum concentration was calculated from the barcode counts of each pseudovirus variant relative to the non-neutralized standard. Then the *polyclonal* (*81*) software (v6.11) was used to fit neutralization curves and estimates of mutation effects on neutralization to these variant-level fraction infectivity measurements. An example notebook for data fitting for mouse sera is at https://dms-vep.org/Flu_H5_American-Wigeon_South-Carolina_2021-H5N1_DMS/notebooks/avg_escape_antibody_escape_mouse-1-03.html and for ferret data is at https://dms-vep.org/Flu_H5_American-Wigeon_South-Carolina_2021-H5N1_DMS/notebooks/avg_escape_antibody_escape_ferret-7.html.

### H5N1 lentiviral pseudovirus for neutralization assays

To determine sera neutralizing concentrations needed to perform the deep mutational scanning experiments to measure the effects of mutations on neutralization (described in the previous methods subsection), lentiviral pseudovirus neutralization assays were used. H5N1 pseudoviruses were produced produced by transfecting 8e5 293T cells plated in wells of a 6-well plate with luciferase and zsGreen coding lentivirus backbone (1 µg per well, plasmid map https://github.com/dms-vep/Flu_H5_American-Wigeon_South-Carolina_2021-H5N1_DMS/blob/main/library_design/2727_pHAGE6-wtCMV-Luc2-BrCr1-ZsGreen-W-1247.gb), lentiviral Gag/Pol, Tat and Rev helper plasmids (0.25 µg per well each), neuraminidase expression plasmid (0.05 µg per well), and a H5 HA expression plasmid (0.25 µg per well, plasmid map https://github.com/dms-vep/Flu_H5_American-Wigeon_South-Carolina_2021-H5N1_DMS/blob/main/library_design/HDM_H5_American_Wigeon.gb). 48 hours after transfection cell supernatants were collected, filtered through a surfactant-free cellulose acetate 0.45 μm syringe filter (Corning, Cat. No. 431220) and stored at −80 °C until further use. Pseudovirus titers were determined by serially diluting virus stocks on 293T cells in the presence of 1 µM oseltamivir and 2.5 µg/ml amphotericin B and measuring luciferase signal 48 hours after infection using Bright-Glo Luciferase Assay System (Promega, E2610), as described previously (*83*).

To perform neutralization assays 2.5e4 293T cells were plated per well in poly-L-lysine coated, black-walled, 96-well plates (Greiner 655930) in the presence of 2.5 µg/ml amphotericin B. The next day sera, were serially diluted in D10 media and incubated with H5N1 pseudovirus in the presence of 1 µM oseltamivir for 45 min at 37 °C. After incubation virus and sera dilutions were transferred onto 293T cells. At 48 hours after infection Bright-Glo Luciferase Assay System (Promega, E2610) was used to measure virus neutralization. The *neutcurve* software (*84*) (https://jbloomlab.github.io/neutcurve/) was used to fit Hill curves and plot neutralization curves from which the appropriate IC99 values were estimated.

### Production of conditionally replicative influenza viruses

Conditionally replicative influenza viruses with wild-type, mutant or selected strain HA genes were generated as schematized in **fig. S2C** and described previously (*85*, *86*) with several modifications explained here. The native HA sequences (not codon optimized) lacking the polybasic cleavage site for all strains and variants were cloned into the bidirectional pHW2000 influenza reverse genetics plasmid (*87*) (plasmid maps for all strains can be found at https://github.com/dms-vep/Flu_H5_American-Wigeon_South-Carolina_2021-H5N1_DMS/tree/main/library_design/conditionally_replicative_virus_plasmids). The polybasic cleavage site was removed from the HA sequence to further enhance the safety of this system. The native NA sequence from A/turkey/Indiana/22-003707-003/2022 strain was also cloned into a pHW2000 plasmid (plasmid map https://github.com/dms-vep/Flu_H5_American-Wigeon_South-Carolina_2021-H5N1_DMS/blob/main/library_design/conditionally_replicative_virus_plasmids/3972_pHW_N1_TurkeyH5.gb). A mixture of 5e5 293T-PB1 cells (*29*) and 50,000 MDCK-SIAT1-PB1-TMPRSS2 cells (*88*) per well was plated into a 6-well plate. The next day cells were transfected with plasmid mix consisting of A/WSN/1933 PB2, PA, NP, M and NS reverse genetics plasmids (*87*), pHH-PB1flank-eGFP plasmid that consists of non-coding terminal regions from A/WSN/1933 PB1 segment and eGFP gene in place of PB1 coding sequence (*29*), pHW_HA_American-Wigeon_USDA-000345-001_H5N1_delPBCS plasmid and pHW_N1_TurkeyH5 plasmid. Next day cell media was changed to Influenza growth media (see Cell lines methods subsection below) and virus supernatant was collected 48 hours after media change. Virus titers were determined by serially diluting virus stocks and infecting MDCK-SIAT1-PB1 cells in Neutralization Assay Medium (NAM) media. 16 hours post infection flow cytometry was used to determine eGFP positive cell numbers and Poisson distribution was used to determine virus titers. In cases where high virus titers were needed, rescued viruses were further passaged at low MOI (0.01) in MDCK-SIAT1-PB1-TMPRSS2 cells in NAM. For stability mutation validations, passaged viruses were further concentrated using 100k Amicon spin columns. To avoid any possibility of reassortment between the conditionally replicative influenza viruses with the H5N1 HA and NA and fully replication competent influenza, work with these conditionally replicative viruses was strictly segregated away from tissue-culture incubators and biosafety cabinets where fully replicative seasonal or lab-adapted influenza viruses were being studied.

### Validation of stability enhancing mutations using conditionally replicative influenza virus

MDCK-SIAT1-PB1 cells were plated in NAM at 100 k per well in a 6-well dish, 4 hours prior to infection. Wild-type and mutant viruses were diluted to the target final MOI of 0.75-1 and incubated with citrate buffers at pHs 6.9, 5.7, 5.5, or 5.3 for 30 minutes at 37°C at a dilution of 4 µl virus to 96 µl of buffer. Citrate buffers were prepared at 0.1 M by mixing together sodium citrate dihydrate and citric acid solutions and adjusted to the required pH using hydrochloric acid or sodium hydroxide. pH treated viruses were then brought up to a neutral pH by 3.9mL of NAM media. 2mL of the NAM diluted virus was then added to MDCK-SIAT1-PB1 cells. At 16-20 hours post infection, cells were collected for flow cytometry analysis and the percent positive eGFP cells were measured for each mutant. We conducted single replicates of each mutant and repeated these across 3 different days. The fractional infectivity was then calculated by taking the proportion of the percent positive green cells at pH 5.3 or pH 5.5 over the percent positive green cells at pH 6.9 (**fig. S5**).

### Neutralization assays using conditionally replicative influenza virus

To validate sera escape mutations we performed neutralization assays using conditionally replicative virus as described previously (*85*). In brief, 50 k MDCK-SIAT1-PB1 were plated into 96-well plates in NAM media 4 hours before infection. Sera were serially diluted in NAM media and mixed with wild-type or mutant conditionally replicative influenza viruses. The amount of virus to use was chosen such that the green fluorescence signal would change linearly with degree of sera neutralization. Virus and sera were incubated for 45 min at 37 °C and pre-plated MDCK-SIAT1-PB1 were infected. 16 hours post infection fluorescence signal was determined using Tecan M1000 plate reader. Fraction infectivity was determined relative to no serum controls and neutralization curves were plotted using *neutcurve* software (*84*).

### Phylogenetic analysis

All available H5Nx HA sequences were downloaded from GISAID (gisaid.org); an acknowledgment list including accession numbers, originating labs, and submitting labs for all GISAID sequences used for phylogenetic analyses is provided in **Supplemental Information 1**. To visualize the DMS results in the context of H5Nx phylogenies, three trees were built: one that included a sample of historical H5Nx HA sequences (all clades), one that included clade 2.3.4.4b HA sequences, and that was subsetted to clade 2.3.4.4b sequences sampled recently in the United States. All phylogenies were constructed with HA sequences only. For the H5Nx all clades tree, sequences were randomly subsampled to 2 sequences per host per country per clade per month. For the clade 2.3.4.4b trees, sequences were randomly subsampled to 4 sequences per country per month (all 2.3.4.4b), or 25 sequences per month (United States). To maximize the ability to identify circulating sequences harboring putative adaptive mutations, all mammalian sequences were force-included in each build. Only HA sequences of at least 1650 nucleotides were included. Sequences were excluded if the isolate was environmental or laboratory-derived or if it was collected before 1996 (H5Nx), 2009 (clade 2.3.4.4b), or 2022 (United States). Using the Nextstrain pipeline (*60*), sequences were aligned with MAFFT (*88*), divergence phylogenies were created using IQ-TREE 2 (*90*), and time-resolved phylogenies were generated with TreeTime (*91*) with ancestral state reconstruction of nucleotide and amino acid sequences. DMS phenotypes were then mapped onto trees by summing the effect values of HA substitutions relative to the DMS strain, A/American Wigeon/South Carolina/USDA-000345-001/2021, for both terminal tips and reconstructed internal nodes (code provided at https://github.com/moncla-lab/annotate-dms). For mouse and ferret sera escape, only positive effect values were summed, while for HA stability, cell entry, and α2-6 sialic acid usage both positive and negative effect values were used.

### Serum generation

Murine experiments were approved by the Institutional Animal Care and Use Committees (IACUC) of the Wistar Institute and the University of Pennsylvania. Ten female C57BL/6 mice (Charles River Laboratories) aged 6-8 weeks were primed intramuscularly with a combination of 5 μg clade 2.3.4.4b H5 rHA + AddaVax and boosted 28 days later with the same vaccine. For immunizations, equal volumes of recombinant protein and AddaVax were mixed according to manufacturer’s instructions. Blood samples were obtained by terminal bleeding 28 days after boost and pooled sera were isolated by centrifugation using Z-Gel tubes (Sarsedt).

Ferret experiments were approved by St. Jude Children’s Research Hospital Institutional Animal Care and Use Committee (IACUC, protocol number 428) according to the guidelines established by the Institute of Laboratory Animal Resources, approved by the Governing Board of the US National Research Council, and executed by trained personnel in a United States Department of Agriculture (USDA)-inspected Animal Biosafety Level 3+ animal facility according to regulations established by the Division of Agricultural Select Agents and Toxins at the USDA Animal and Plant Health Inspection Service (APHIS), as governed by the United States Federal Select Agent Program (FSAP) regulations (7 CFR Part 331, 9 CFR Part 121.3, 42 CFR Part 73.3). For ferret infections male ferrets (Triple F Farms, Sayre, PA, USA) were used. Ferrets 10 and 19 were intranasally inoculated at 10^6 TCID50s and terminal bleeds were collected 25 and 28 days post infection, respectively. Ferret 7 was intramuscularly primed with inactivated whole virus and 14 days later challenged intranasally with 10^3 TCID50s. H5 strains used for ferret infections are indicated in **fig. S6**. The ferret sera used in this study was all residual sera from prior experiments.

All sera were inactivated for 1h at 56°C to eliminate complement activity and ferret sera were treated with receptor-destroying enzyme.

### Cell lines and media

The HEK293 isogenic α2-3 cells and α2-6 cells were stably engineered as described previously (*31*, *92*). α2-3 cells are 293 cells with combinatorial knockout of ST3GAL3, ST3GAL4, ST3GAL6, ST6GAL1 and ST6GAL2 genes and knockin of the human ST3GAL4 gene. α2-6 cells are 293 cells with knockout of ST3GAL3, ST3GAL4, ST3GAL6, ST6GAL1 and ST6GAL2 genes and knockin of the human ST6GAL1 gene. Note that both cell lines still express α2-3 and α2-6-linked sialic acids on O-linked glycans.

These cells and 293T cells were maintained D10 media (Dulbecco’s Modified Eagle Medium with 10% heat-inactivated fetal bovine serum, 2 mM l-glutamine, 100 U/mL penicillin, and 100 μg/mL streptomycin). D10 media was used for all experiments involving pseudovirus. MDCK-SIAT1-PB1 and MDCK-SIAT1-PB1-TMPRSS2 cells were passaged in D10 media and for assays described above plated in either Influenza growth media (Opti-MEM supplemented with 0.1% heat-inactivated FBS, 0.3% bovine serum albumin, 100 µg/mL of calcium chloride, 100 U/mL penicillin, and 100 µg/mL streptomycin), or Neutralization assay media (Medium-199 supplemented with 0.01% heat-inactivated FBS, 0.3% BSA, 100 U/mL penicillin, 100 μg/mL streptomycin, 100 μg/mL calcium chloride, and 25mM HEPES). Influenza growth media was used for all conditionally replicative virus rescue experiments, while Neutralization assay media was used for all experiments involving fluorescence readings because riboflavin present in Opti-MEM has high background fluorescence.

---

**Figure S1.**
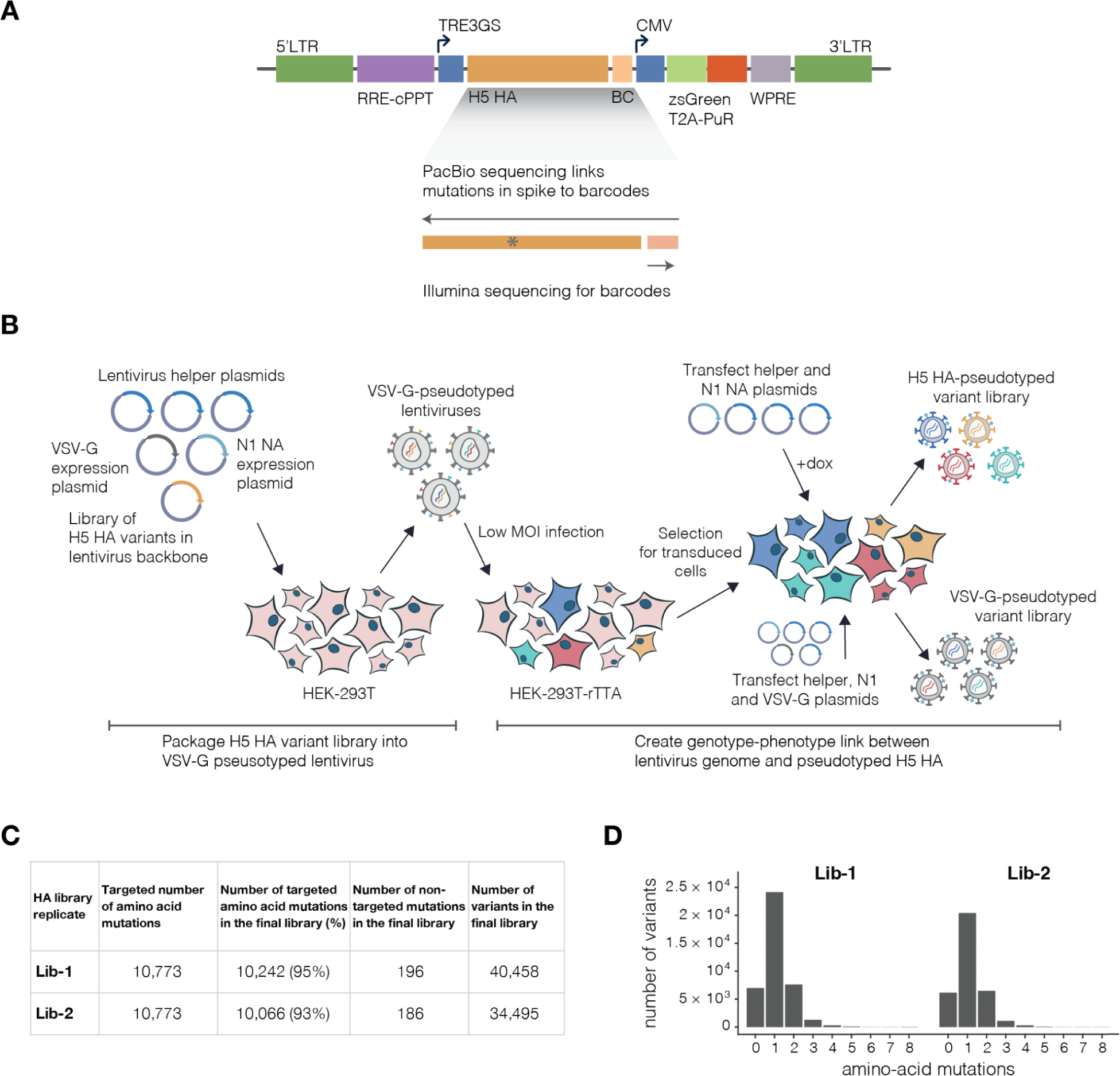
H5 HA deep mutational scanning libraries. **A**, Schematic of lentiviral backbone used to produce the pseudoviruses. The HA gene is encoded in the lentiviral backbone followed by a 16-nucleotide barcode after the stop codon. Other notable elements include the 5’ LTR, an inducible TRE3GS promoter that drives HA expression, a constitutively expressed ZsGreen / puromycin cassette, and a repaired 3’ LTR that allows re-activation of integrated proviral genomes. The full sequence of each HA mutant is linked to the barcode by long-read PacBio sequencing, enabling subsequent experiments to simply sequence the short barcode to identify the full HA mutant. Note that this backbone has previously been described (*18*), and the full plasmid encoding mCherry rather than HA is available on AddGene as item 204579. **B**, A two-step process is used to create genotype-phenotype linked viruses pseudotyped with HA, as described previously for other viral entry proteins (*18*). First, VSV-G pseudotyped lentiviral particles are generated by standard transfection, including a plasmid expressing the influenza NA gene to ensure virions do not remain attached to cells via the HA protein. These lentiviral particles are then used to infect 293T-rTTA cells at low multiplicity of infection such that infected cells typically just have one integrated provirus, and cells containing proviruses are then selected using puromycin. Because the lentiviral backbone contains a full 3’ LTR, the proviral genomes can be re-activated by transfection of the lentiviral helper plasmids and a NA expression plasmid to produce a library of genotype-phenotype linked HA-pseudotyped virions. For selections designed to measure HA cell entry, we also produce virions pseudotyped with VSV-G to use as a control comparison, with cell entry quantified by the relative infectivity of each HA variant in the presence versus absence of VSV-G (since the VSV-G pseudotyped particles do not require a functional HA for cell entry; see **fig. S2A**). **C**, Number of targeted and actual HA amino-acid mutations found in each of the two replicate mutant libraries. The last column indicates the number of unique barcoded HAs in each library; this number exceeds the number of unique mutations as many mutations are found in multiple different barcoded HA variants. **D**, Number of HA amino-acid mutations in each barcoded HA variant in each library.

**Figure S2.**
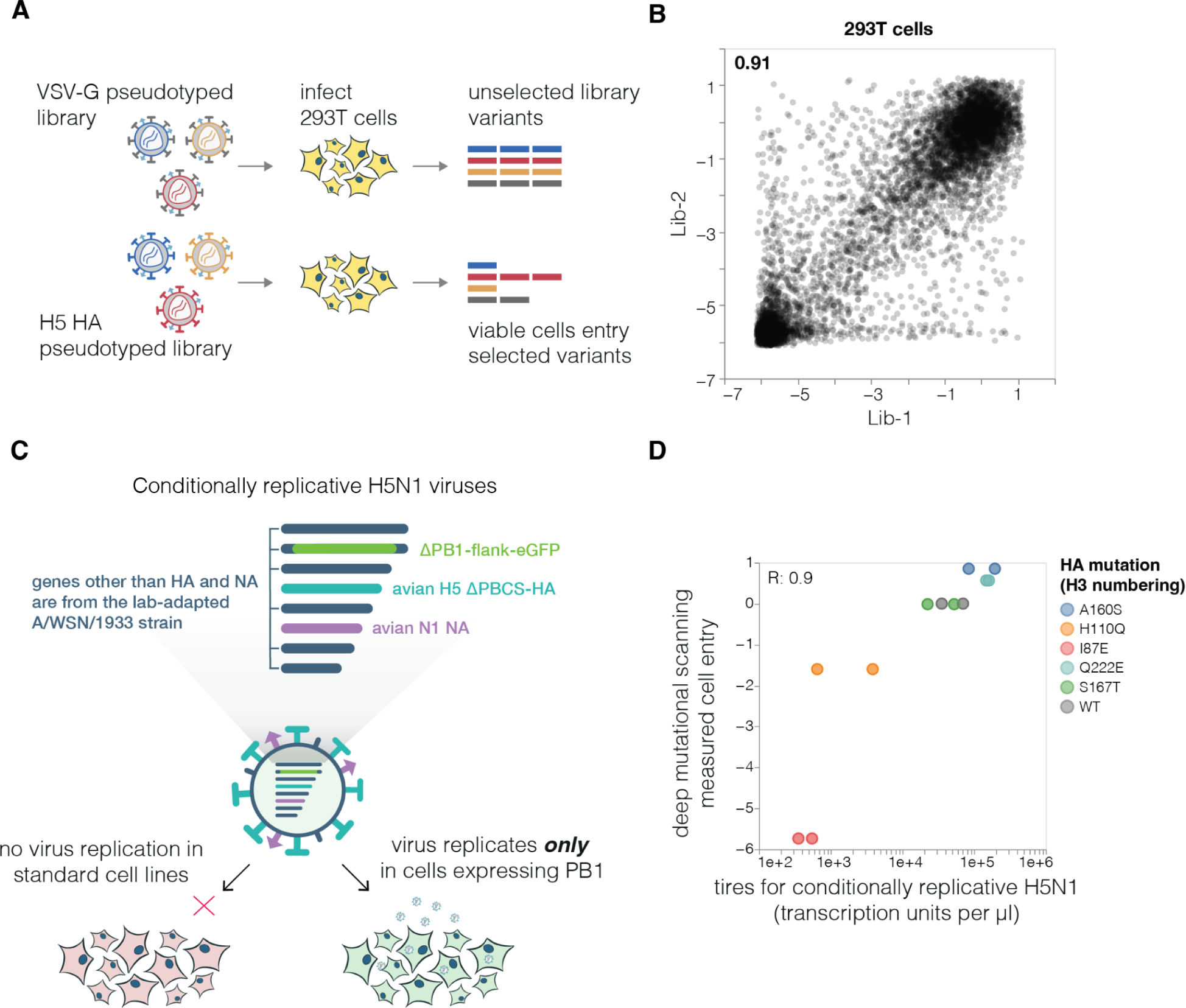
Deep mutational scanning for cell entry and validation using conditionally replicative influenza virions. **A**, The effects of mutations on cell entry were measured by generating the pseudovirus libraries with virions expressing either the mutant HAs or unmodified VSV-G on their surface (see **fig. S1B**). The ability of each barcoded variant to enter 293T cells was then compared between the VSV-G and HA pseudotyped libraries. Entry of the HA pseudotyped variants depends on the function of HA, whereas entry of the VSV-G pseudotyped variants is independent of HA function. Therefore, the effects of HA mutations on cell entry can be determined by comparing the frequency of each variant between the two conditions. **B**, Correlation between mutational effects on cell entry in 293T cells as measured in each of the two independent replicate libraries. The Pearson correlation is indicated at top left. Throughout the rest of this paper, we report the average effect across the two library replicates. **C**, For validation assays, we used conditionally replicative influenza virions that can be safely studied at biosafety-level 2. The virions have the essential PB1 polymerase gene replaced by eGFP (*29*) and derive their non-HA/NA proteins from the lab-adapted A/WSN/1933 (H1N1) strain (*30*). These viruses can be generated by reverse genetics and propagated in cells that constitutively express the PB1 protein, but cannot undergo replication in standard cells as they do not express PB1. **D**, Deep mutational scanning measurements of how the indicated mutations affect cell entry in 293T cells versus the titers of conditionally replicative influenza virions carrying the same mutations on MDCK-SIAT1-TMPRSS2-PB1 and 293T-PB1 co-cultures. The titers of the conditionally replicative influenza virions represent those in the supernatant at 48 hours after generating the virions by reverse genetics. There are two points for each mutation for the conditionally replicative influenza virions because viruses were rescued twice from independent plasmid preparations. The mutations selected for validation were chosen to span a range of cell-entry effects. R indicates the Pearson correlation.

**Figure S3.**
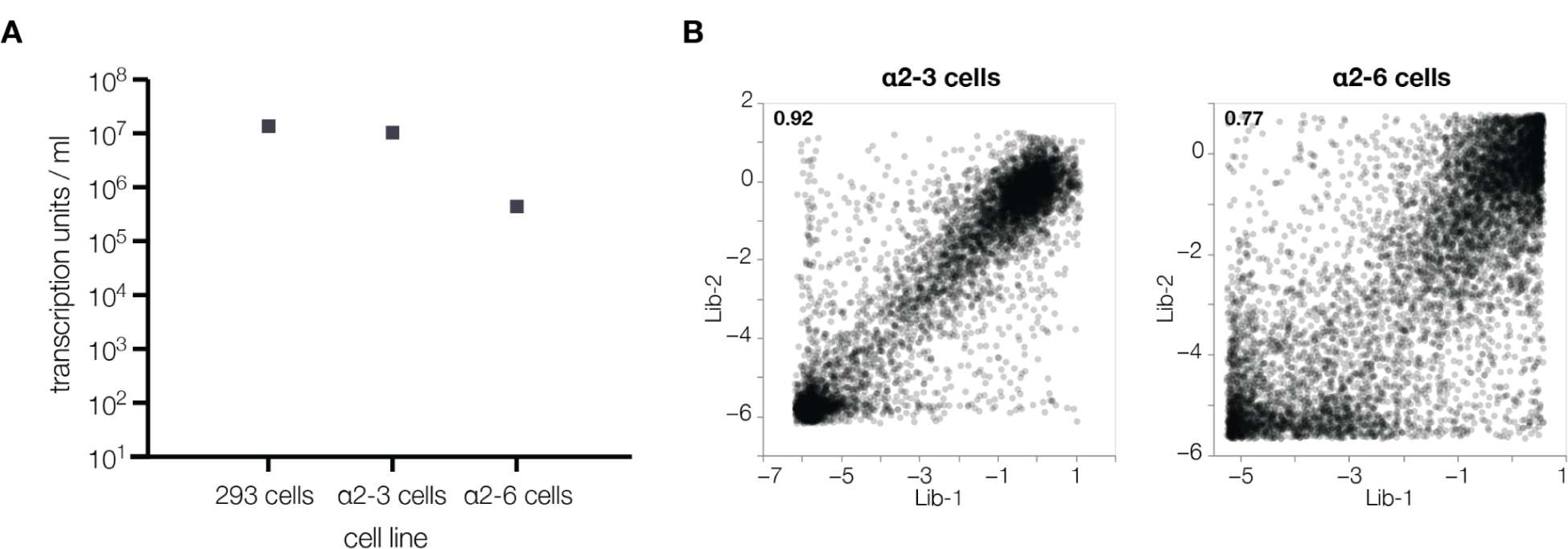
H5 pseudovirus titers on cell lines expressing different sialic-acid receptors. **A**, Titers of the lentiviral particles pseudotyped with the unmutated H5 HA on standard 293 cells and the α2-3 or α2-6 293 cells that express α2-3-linked or α2-6-linked sialic acids (*31*). The virions can infect all cells, but the titers are substantially lower on α2-6 cells, consistent with the fact that avian H5 HAs are better at utilizing α2-3-linked sialic acids. **B**, Correlation between the effects of mutation on entry in α2-3 and α2-6 cells as measured with each of the two independent replicate pseudovirus libraries. The Pearson correlation is indicated at the top left. The measurements are noisier (more poorly correlated between replicates) for the α2-6 cells as there is more noise due to worse entry into those cells. Throughout the rest of the paper, we report the average effect across the two library replicates.

**Figure S4.**
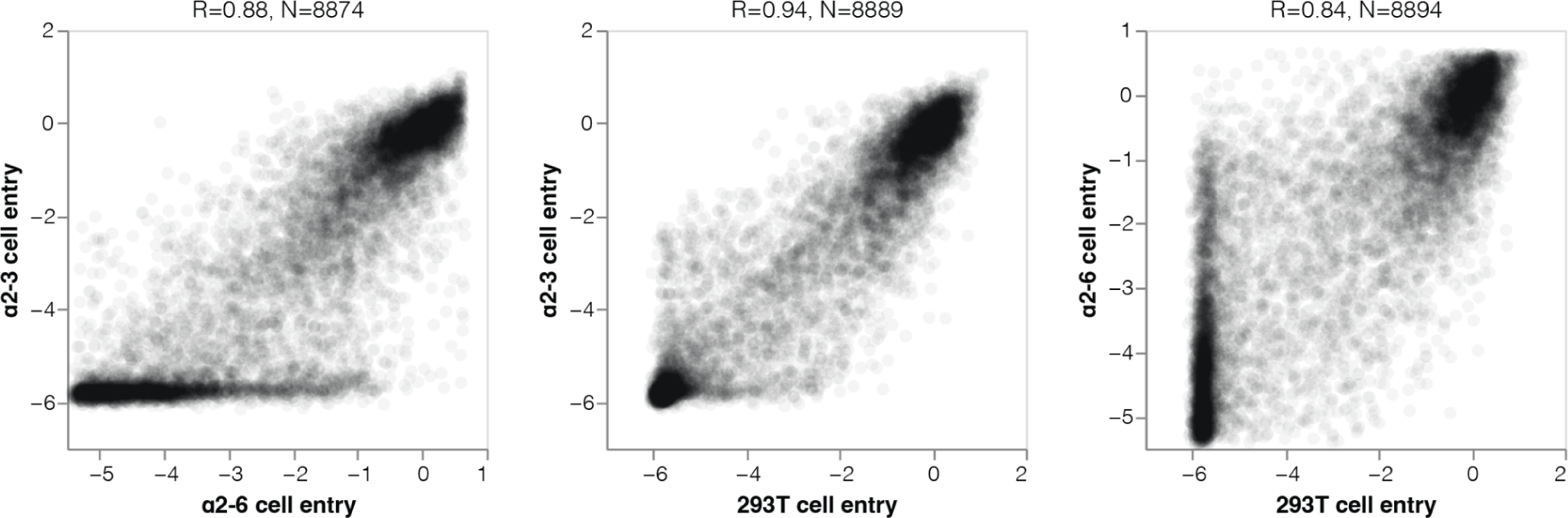
Correlations among effects of HA mutations on entry into different cell lines. Correlations among the measured effects of mutations on entry into 293T, α2-3, and α2-6 cells. Note that the measurements for entry into α2-6 cells are more poorly correlated with the other cells for two reasons. First, there are some mutations that actually increase usage of α2-6-linked sialic acid as highlighted in Fig. 3; these mutations are characterized by good overall entry into the α2-6 cells relative to other cells. But additionally, since the parental H5 HA used in our experiments is relatively poor at entering the α2-6 cells and mutation effects are relative to that parent, many mutations that are poor overall for cell entry nonetheless are less deleterious relative the parent (not as negative cell entry effect) on α2-6 cells compared to the other cells. In the scatter plots, R indicates the Pearson correlation. See https://dms-vep.org/Flu_H5_American-Wigeon_South-Carolina_2021-H5N1_DMS/a26_usage.html for more detailed interactive plots of the effects of mutations on entry into the different cells.

**Figure S5.**
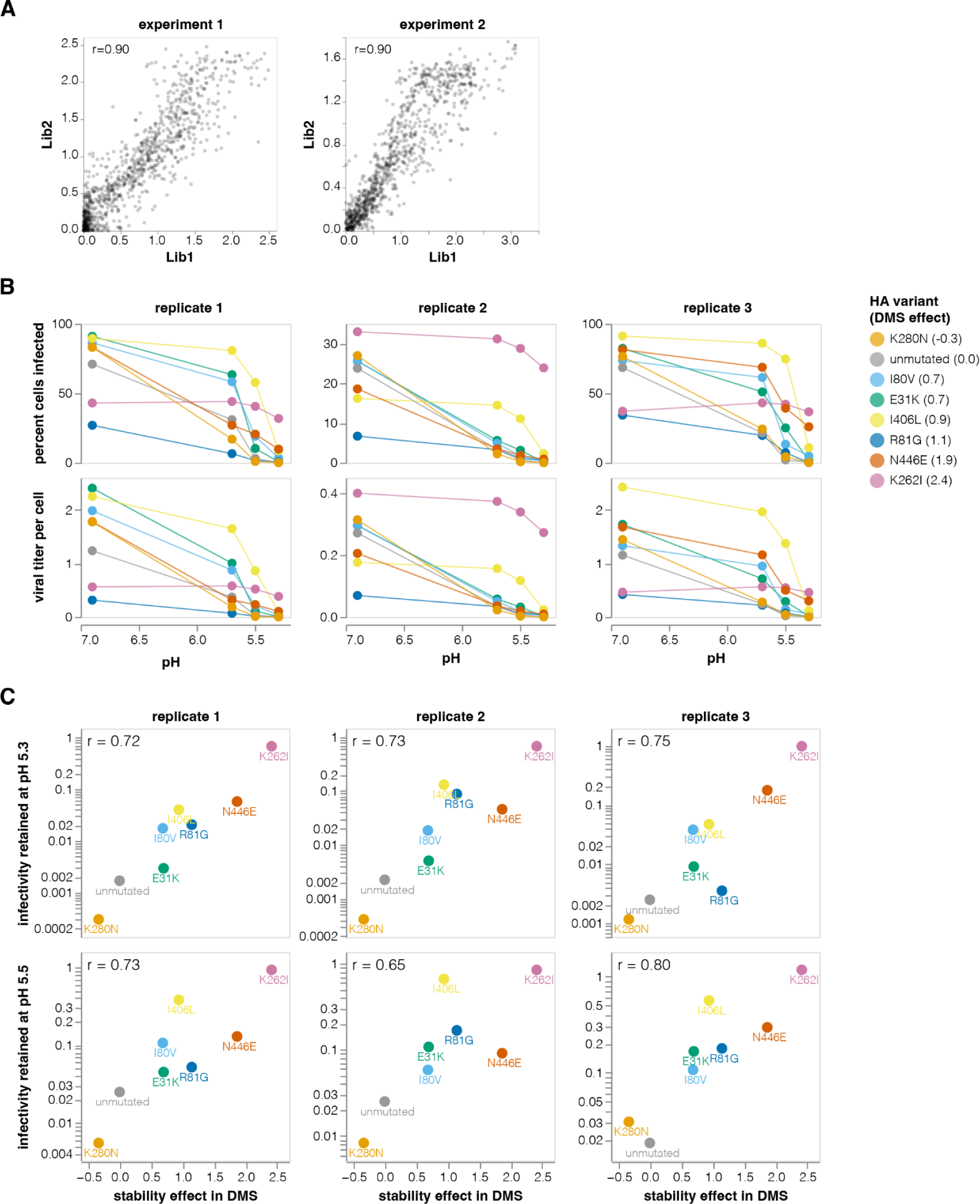
Validation of stability enhancing HA mutations. **A**, Correlations between the effects of mutations on HA stability as measured by the deep mutational scanning for each of the two independent library replicates. Experiment 1 and 2 were performed on different dates. Only stabilizing mutations are shown. The Pearson correlation is shown in the upper left. Throughout the rest of this paper we report the average effect across libraries and experiments. **B-C,** Stability-enhancing mutations identified in deep mutational scanning were validated using the conditionally replicative influenza virions. **B**, Conditionally replicative influenza virions carrying each of the indicated mutations were incubated at a buffer of the indicated pH for 30 minutes, diluted into a neutral pH buffer, and then used to infect target cells. The top row shows the percent of cells infected by each virus after treatment at each pH, and the bottom row shows the titer per cell (MOI) calculated using the Poisson distribution. **C**, The infectivity remaining for each mutant at the indicated pHs relative to pH 6.9 (as calculated from the bottom row of panel **B**) versus the stability effects of the mutation measured in the deep mutational scanning. The Pearson correlation is indicated in the upper-left of each plot.

**Figure S6.**
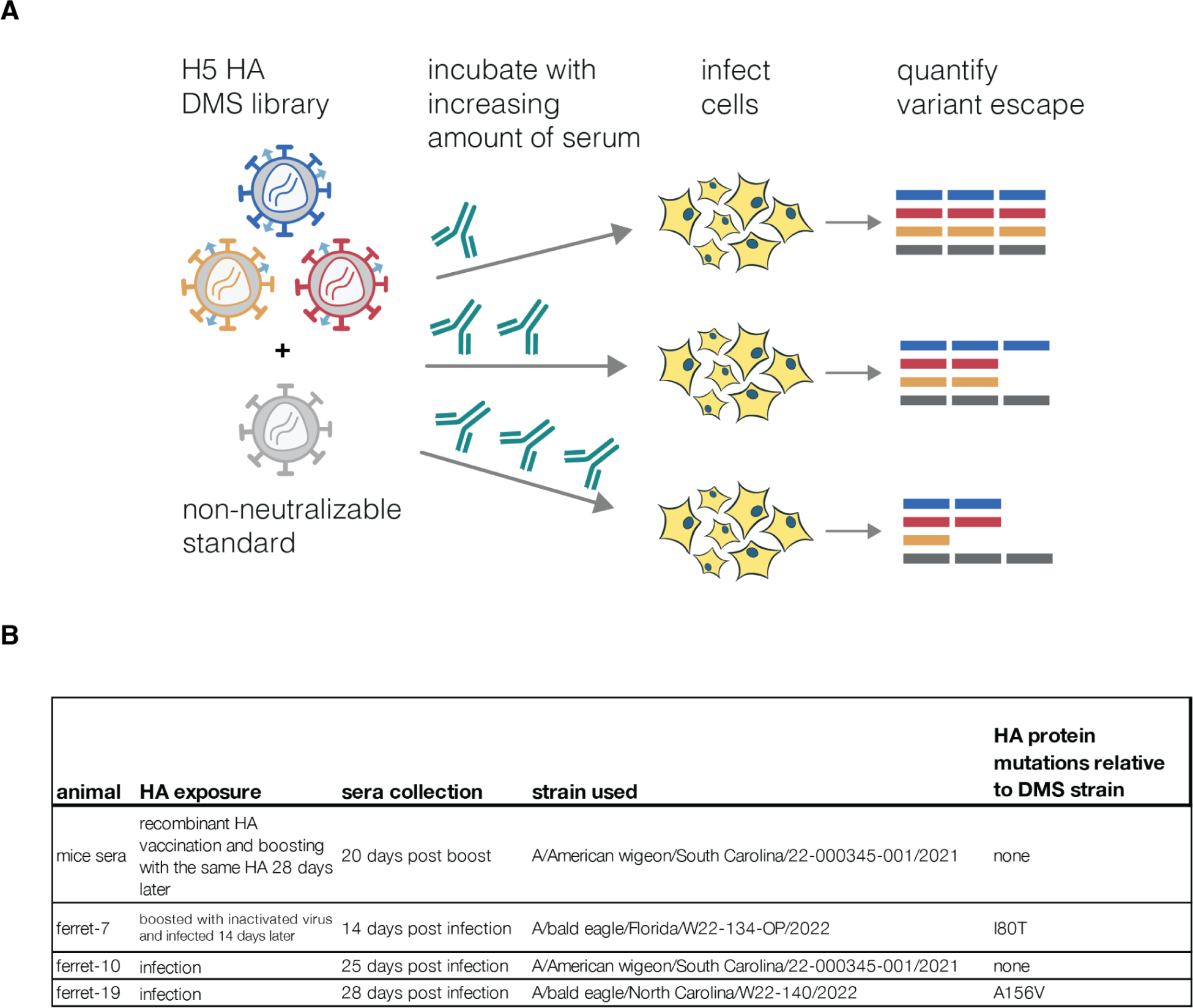
Measuring escape from sera neutralization. **A**, Workflow for measuring escape from polyclonal serum using the pseudovirus libraries. The libraries are combined with a non-neutralizable standard virus that is not affected by the serum antibodies (in this case VSV-G pseudovirus was used for mice sera and RdPro pseudovirus for ferret sera), and then incubated with increasing concentrations of sera and infected into 293T cells. Neutralization is quantified by comparing the frequency of each viral mutant barcode to that of the non-neutralizable standard, as described previously (*18*). **B**, All mice sera came from mice that were vaccinated and boosted with recombinant HA protein that exactly matched the parental strain used in the deep mutational scanning. Ferret sera came from animals that were infected with 2.3.4.4b clade viruses or infected and then boosted with matched inactivated virus. For two of the ferrets, the immunizing strain differed by a single amino-acid mutation from the parental strain used in the deep mutational scanning.

**Figure S7.**
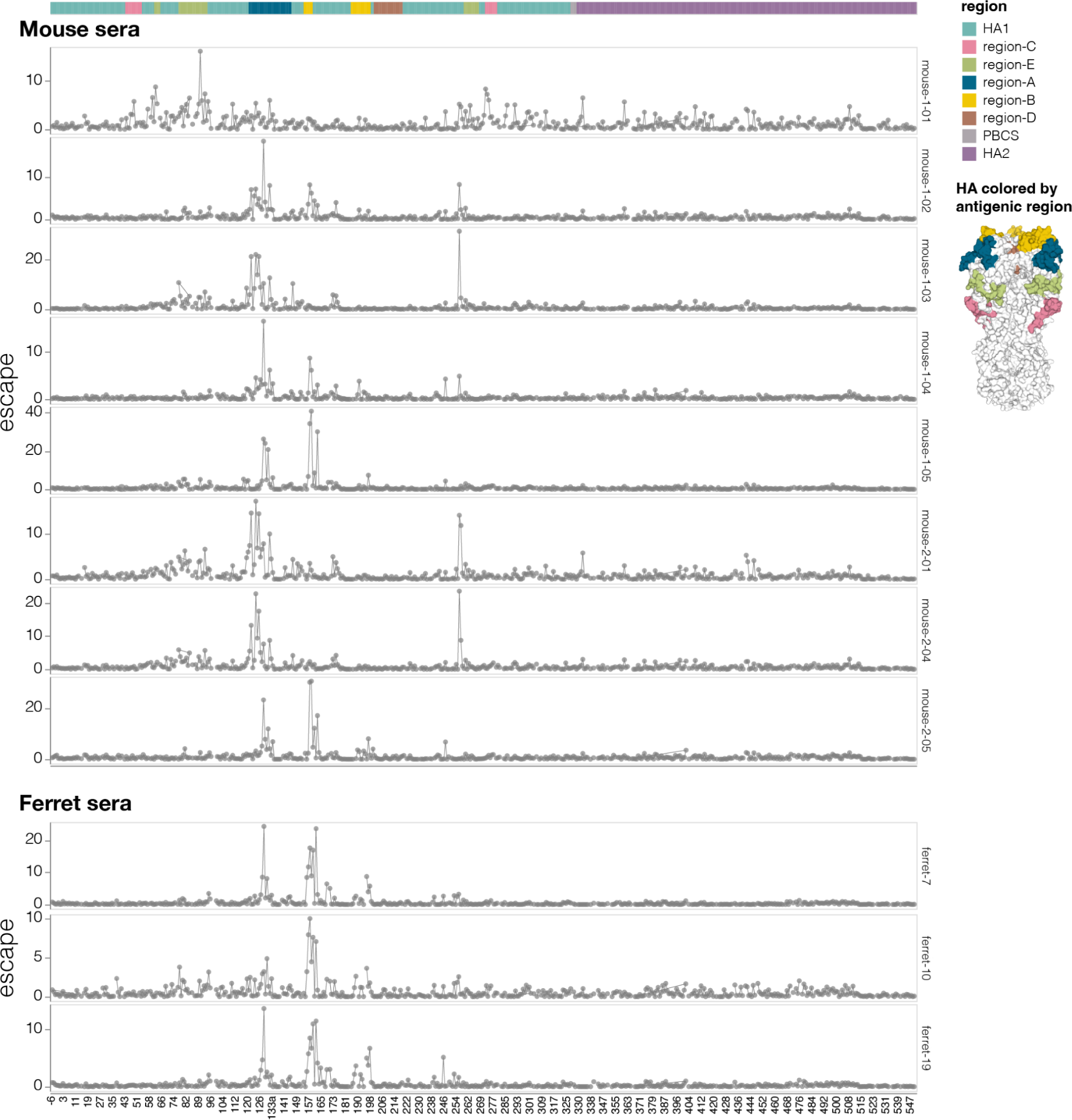
Effects of mutations on escape from neutralization for sera from individual mice and ferrets. Total escape caused by mutations at each site in HA for individual mice and ferret sera used in this study. There is some animal-to-animal variation even within species: for instance, the sera of most mice is primarily escaped by mutations in antigenic regions A and B, but the serum from mouse-1-01 is primarily escaped by mutations in antigenic region E. See https://dms-vep.org/Flu_H5_American-Wigeon_South-Carolina_2021-H5N1_DMS/escape.html for interactive plots.

**Figure S8.**
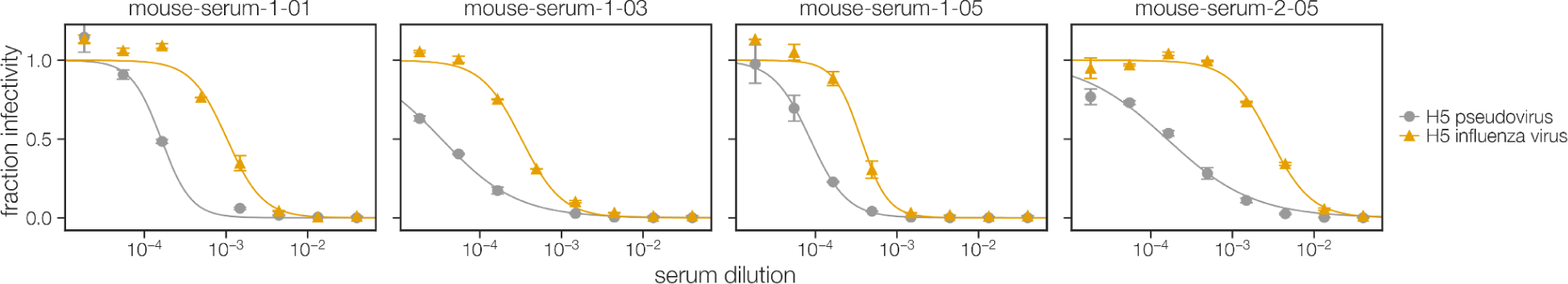
Neutralization of pseudoviruses versus conditionally replicative influenza viruses. Comparison between mouse sera neutralization of A/AmericanWigeon/SouthCarolina/USDA-000345-001/2021 pseudovirus and conditionally replicative influenza virus with the same HA. Note that the pseudoviruses (pseudotyped lentiviral particles) are more sensitive to neutralization than the actual influenza virions.

**Figure S9.**
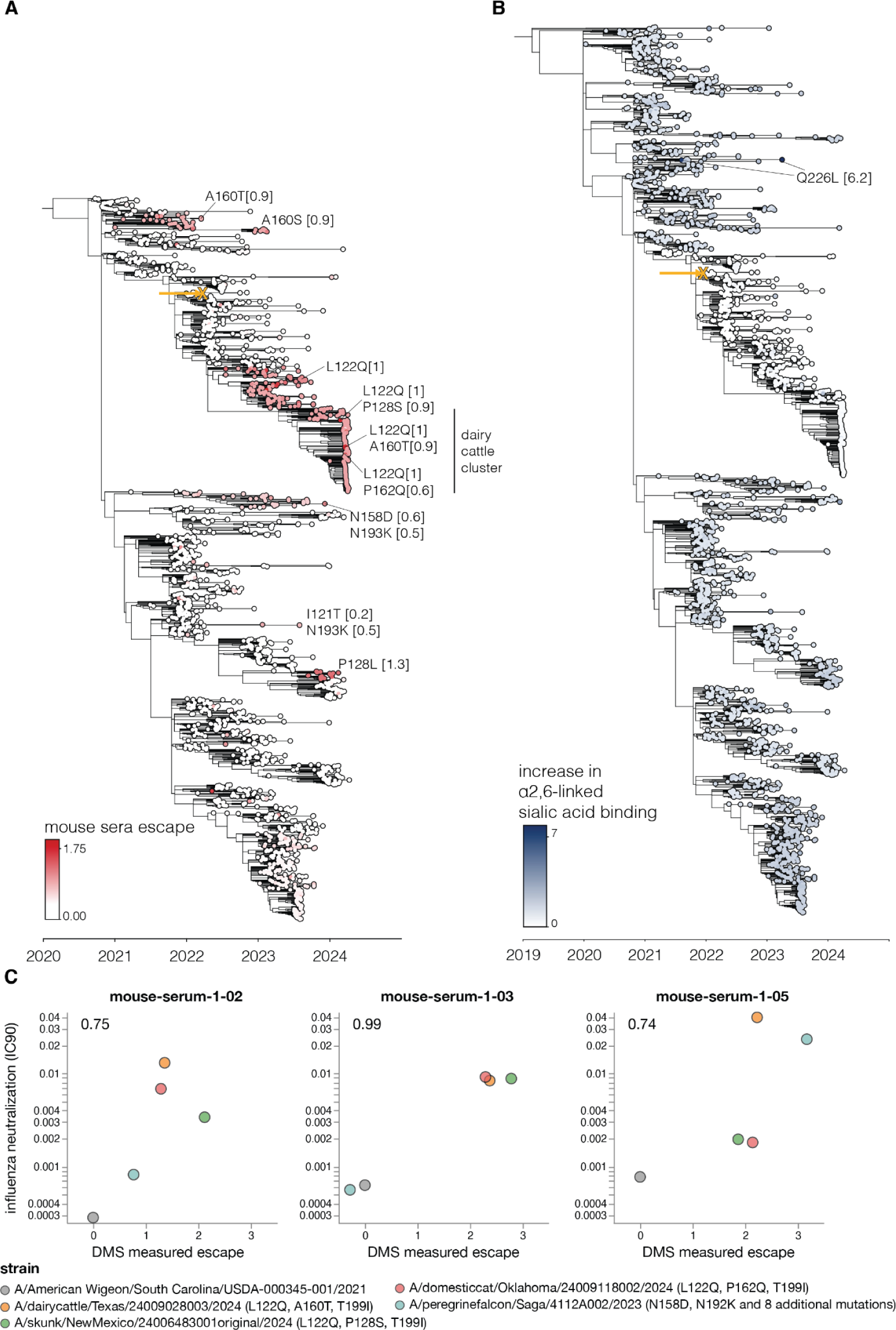
Mouse sera neutralization and α2-6-linked sialic acid binding increase for circulating 2.3.4.4b strains. Trees similar to those in Fig. 6 but showing different phenotypes, namely (**A**) mouse sera escape and (**B**) increase in α2-6-linked sialic acid binding. **B** shows sequences starting from 2019 whereas **A** only shows sequences starting from 2020. **C** HAs for selected strains carrying mutations that caused strong escape from sera neutralization in the deep mutational scanning were assayed for neutralization by the mouse sera in influenza-based neutralization assays (**fig. S10B**). These scatter plots show the neutralization measured in the influenza virus assays (y-axis) versus the predicted escape calculated as the simple sum of the mutations in each HA relative to the strain used in the deep mutational scanning. The full list of mutations in each strain is provided in **fig. S10D**.

**Figure S10.**
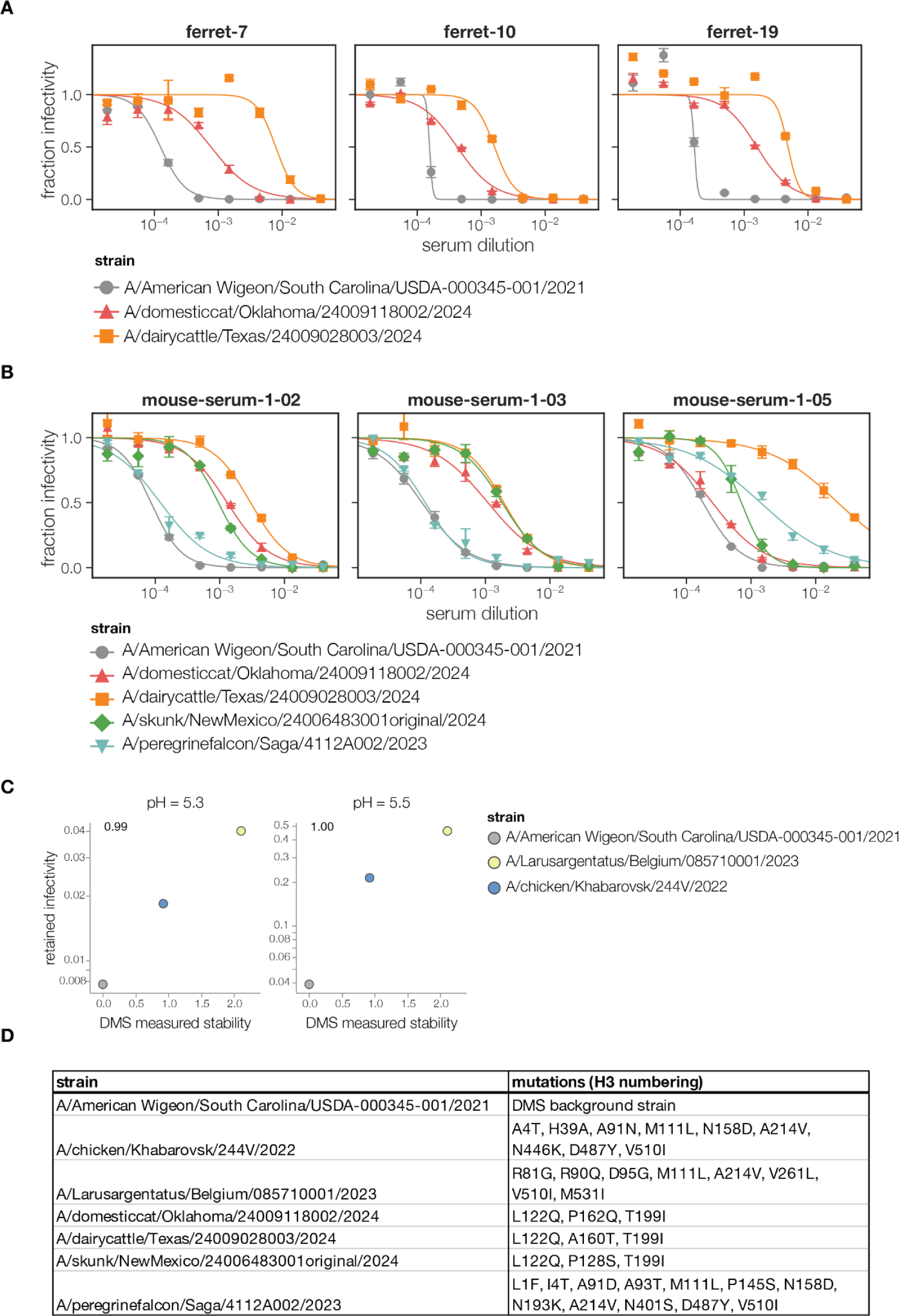
Sera neutralization and stability of circulating 2.3.4.4b strains. **A** Ferret and **B** mouse sera neutralization curves for 2.3.4.4b clade strains measured using conditionally replicative influenza virus These are the neutralization curves that generated the data plotted in Fig. 6C and **fig. S9C**. **C** same as Fig. 6D but with infectivity measured at pH 5.3 and 5.5. **D** A list of HA amino-acid mutations (H3 numbering) in circulating 2.3.4.4b strains used for validation relative to the A/AmericanWigeon/SouthCarolina/USDA-000345-001/2021 strain used for the deep mutational scanning.

